# Presynapses are mitophagy pit stops that prevent axon degeneration

**DOI:** 10.1101/2024.09.09.611943

**Authors:** Wai Kit Lam, Runa S. J. Lindblom, Bridget Milky, Paris Mazzachi, Marjan Hadian-Jazi, Catharina Küng, Grace Khuu, Louise Uoselis, Thanh Ngoc Nguyen, Marvin Skulsuppaisarn, Tahnee L. Saunders, Marlene F. Schmidt, Grant Dewson, Adam I. Fogel, Cedric Bardy, Michael Lazarou

**Author notes:** Department of Biochemistry and Molecular Biology, Biomedicine Discovery Institute, Monash University, Melbourne, Australia (Marjan Hadian-Jazi).

## Abstract

Defects in neuronal mitophagy have been linked to neurodegenerative diseases including Parkinson’s disease. However, despite the importance of mitophagy in neuronal homeostasis, the mechanistic basis for neurodegeneration when mitophagy is defective is unclear. Here, using human neurons, we discover that presynapses are mitophagy pit stops for damaged axonal mitochondria. We show that while mitochondrial damage and PINK1/Parkin activation events are distributed throughout axons, mitophagy initiation and autophagosome formation are restricted to presynapses, which we show contain the machineries required for mitophagy. Being the primary sites of axonal mitophagy, presynapses were vulnerable when PINK1/Parkin mitophagy was defective. We observed local cytochrome *c* release within presynapses from an accumulation of damaged mitochondria. This resulted in downstream degradative caspase activation, defining a mechanism for neurodegeneration. Pharmacological rescue of axon degeneration was achieved through synthetic upregulation of receptor mediated mitophagy with the clinically approved compound Roxadustat, revealing a potential therapeutic avenue for disease.

## Introduction

Mitophagy is a degradative quality control pathway that removes damaged and dysfunctional mitochondria through the macroautophagy system.^1,2^ This involves the encapsulation of damaged mitochondria within a double membrane structure termed an autophagosome that serves as a vehicle to deliver the damaged mitochondria to lysosomes where they are degraded. Autophagosome formation is a tightly orchestrated process that requires the activity of several autophagy machineries, ranging from protein and lipid kinase complexes (ULK1/2, VPS34) to lipid transfer (ATG2A, ATG2B), and lipid scramblase (ATG9A) proteins.^3–10^ A large mobilization of lipid is necessary to build autophagosomes, with the endoplasmic reticulum (ER) being among the key sources of this lipid.^11–13^ The complexity of autophagosome formation creates a conundrum for polarized cells such as neurons in which the necessary machinery must be present at locations distal to the soma, raising the question of what the spatial requirements might be for mitophagy in distal axons.

To date, several different mitophagy pathways have been identified, including PINK1 and Parkin mediated mitophagy, and BNIP3, NIX, or FUNDC1 receptor mediated mitophagy.^14^ PINK1 and Parkin function by generating phospho S65-ubiquitin chains on the surface of damaged mitochondria that recruit cargo adaptors, including Optineurin (OPTN) and NDP52, to initiate mitophagy. PINK1/Parkin mitophagy is largely a mitochondrial damage response pathway. In contrast, mitophagy mediated by receptors such as BNIP3 and NIX is generally thought to drive basal mitophagy, as well as mitophagy in response to cellular differentiation and metabolic rewiring cues.^15,16^ To initiate mitophagy, BNIP3 and NIX directly engage with autophagy machineries including WIPI2 and ATG8 family members.^17–20^ Under steady state conditions, BNIP3 and NIX levels are kept low on mitochondria, at least in part through constitutive ubiquitin dependent turnover by the F-box protein FBXL4 and the mitochondrial phosphatase PPTC7.^21–24^ Transcriptional signalling from HIF1α upregulates BNIP3 and NIX and activates mitophagy.^25–27^

Mitophagy plays an important role in maintaining mitochondrial health, and defects in the pathway have been linked to neurodegenerative diseases, including Parkinson’s disease in which PINK1 and Parkin are mutated.^28,29^ More recently, mitophagy has also been linked to Alzheimer’s disease and amyotrophic lateral sclerosis.^30–32^ However, despite the importance of mitophagy for neuronal health, the mechanisms behind neurodegeneration when mitophagy is defective remain unclear. This includes whether defective mitophagy plays a role in axonal and synaptic vulnerabilities that have been reported in disease including Parkinson’s disease and Alzheimer’s disease.^33–37^ Such work has been hindered, at least in part, because genetic knockout of PINK1 and/or Parkin in rodents has not phenocopied the early onset neurodegeneration of familial Parkinson’s disease,^38–40^ potentially due to developmental compensation.^41^ This has created a roadblock to understanding neurodegenerative mechanisms and highlights the need to analyse the cell biology of mitophagy in human neurons.

Here, using human stem cell derived neurons, we sought to understand the mechanisms of neuronal mitophagy, and explore whether axonal and synaptic vulnerabilities are connected to defective mitophagy. We discover that presynapses serve as axonal mitophagy pit stops that deal with both locally and distally damaged mitochondria. However, we find that this becomes a neuronal vulnerability when PINK1/Parkin mitophagy is defective, as damaged mitochondria accumulate within presynapses where they activate degradative caspases. Pharmacological upregulation of receptor mediated mitophagy via HIF1α signaling, achieved through the clinically used HIF1α prolyl-4 hydroxylase inhibitor Roxadustat, re-activates mitophagy to restore mitochondrial health and protect neurons. Therefore, we propose HIF1α prolyl-4 hydroxylase inhibition as a potential avenue for therapeutic intervention in neurodegenerative diseases with mitophagy impairments.

## Results

### Oxidative stress induces mitophagy in human neurons

To stress mitochondria and analyse mitophagy in human neurons, we used an oxidative stress based approach in which antioxidants are removed from the media^42^. Oxidative damage was chosen since it more closely reflects the type of mitochondrial stress occurring in neurons in physiological settings,^42^ as opposed to pan-mitochondrial uncouplers. First, we established whether antioxidant removal could induce mitochondrial damage and mitophagy using the human i3Neuron system.^43^ In addition, we sought to determine whether this approach damages mitochondria within axons. i3Neurons expressing mito-EBFP2 were stained using the mitochondrial membrane potential dependent dye TMRM (Tetramethylrhodamine, methyl ester). Antioxidant removal for 3-4 h resulted in mitochondrial damage as evidenced by mito-EBFP2 positive mitochondria lacking TMRM staining, demonstrating a loss of membrane potential (Figure 1A). Damaged mitochondria were observed in the soma, in axons, and in axonal swellings/boutons (hereafter boutons). Consistent with the mitochondrial damage induced by antioxidant (AOx) removal, we also observed evidence of mitophagy using imaging of the fluorescent mitophagy reporter mt-Keima at the 5-6 h antioxidant removal time point (Figure 1B). Acidified mt-Keima (552nm) was observed in the soma and axon hillock, but not in axons or axonal boutons, which is consistent with prior observations showing that autophagic lysosomes become acidified upon retrograde transport to the soma.^44–47^

**Figure 1.**
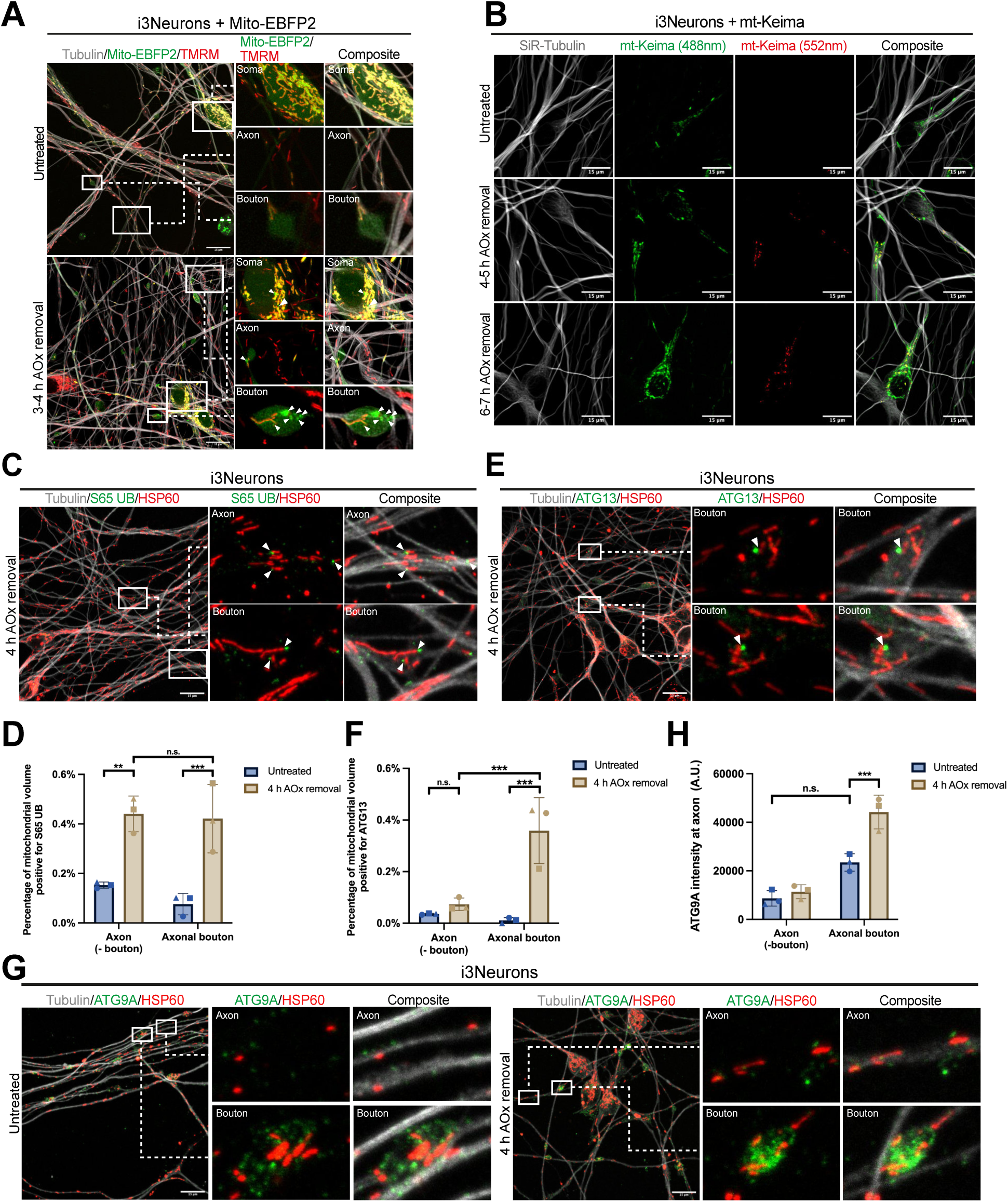
Antioxidant removal damages mitochondria and activates mitophagy in iPSC-derived i3Neurons (A) WT i3Neurons expressing mito-EBFP2 were untreated or cultured in antioxidant (AOx) removal media for 3-4 h and stained with TMRM and SiR-tubulin. White arrowheads indicate mitochondria lacking membrane potential. (B) WT i3Neurons expressing mito-Keima (mt-Keima) were untreated, cultured in AOx removal media for 4-5 h or 6-7 h and stained with SiR-Tubulin. (C and D) WT i3Neurons were untreated or cultured in AOx removal media for 4 h and immunostained for phospho-Serine 65 ubiquitin (S65 UB), HSP60 and tubulin (C) and the percentage of mitochondrial volume positive for S65 UB was quantified (D). S65 UB is a product of PINK1/Parkin activity. White arrowheads point to S65 UB located on mitochondria. (E and F) WT i3Neurons were cultured in AOx removal media for 4 h and immunostained for ATG13, HSP60 and Tubulin (E) and the percentage of mitochondrial volume positive for ATG13 was quantified (F). Untreated samples shown in Figure S1A. White arrowheads indicate ATG13 puncta localised to mitochondria. (G and H) WT i3Neurons were untreated or cultured in AOx removal media for 4 h and immunostained for ATG9A, HSP60 and Tubulin (G) and the ATG9A intensity (Arbitrary unit (A.U.)) at axon was quantified (H). Data in (D), (F) and (H) are mean ± SD from three independent experiments. Two-way ANOVA. **p<0.005, ***p<0.001. n.s., not significant. Scale bars, 15 µm.

### Presynapses are hotspots for axonal mitophagy

Major neurodegenerative disorders, including Parkinson’s disease and Alzheimer’s disease, are thought to begin with axon degeneration. ^33–35^ To explore whether axonal vulnerabilities and mitophagy are linked, we focused on the biology of PINK1/Parkin mitophagy in axons. First, we assessed whether PINK1/Parkin activation and mitophagy initiation occurs within axons, and if so, where. PINK1/Parkin activation events were imaged with phospho-S65-ubiquitin,^48–53^ whereas mitophagy initiation, as defined here by the formation of phagophores on mitochondria, was imaged using ULK1 kinase complex subunit ATG13. A robust increase of phsospho-S65-ubiquitin foci on mitochondria was observed across axons and axonal boutons upon AOx removal (Figures 1C, 1D, and S1A), demonstrating that PINK1/Parkin activation occurs in axons. This is in agreement with previous work showing that PINK1 mRNA is transported along mitochondria in axons.^54^ Having established that PINK1/Parkin activation can occur across axons and axonal boutons using AOx removal, we next assessed ATG13 recruitment to determine where autophagosome formation is initiated in axons. A significant increase of ATG13 foci on mitochondria was observed in axons, but most of these mitophagy initiation events occurred within boutons (Figures 1E, 1F, and S1B), despite the presence of phospho-S65-ubiquitin on mitochondria throughout axons (Figures 1C and 1D). This result shows that while PINK1/Parkin activation can occur along the length of an axon, mitophagy initiation predominantly occurs in boutons. Consistently, the key autophagy factor ATG9A^8^ was found to be enriched in boutons (Figures 1G and 1H), and the signal intensity increased following AOx removal, indicating mitophagy initiation response within the boutons. Taken together, these results demonstrate that mitochondrial damage and PINK1/Parkin activation can occur along the length of axons, whereas mitophagy initiation is largely restricted to boutons.

The enrichment of ATG9A within boutons is indicative of presynaptic sites, as has been shown in *C. elegans* neurons.^55^ We therefore hypothesised that the boutons undergoing mitophagy initiation in i3Neurons represent presynapses. To confirm this, the presence of presynapse markers was assessed using Synapsin I and Synaptophysin (Figures 2A and 2B), which showed enrichment of these markers within boutons, as well as via Western blotting (Figure S2A). Next, we directly assessed whether mitophagy occurs within these presynapse marker enriched boutons. OPTN and NDP52 are the major autophagy adaptors that initiate PINK1/Parkin mitophagy, but unlike NDP52, OPTN is enriched in human brain^56^ and in i3Neurons (Figures 2C and S2B). i3Neurons expressing EGFP-OPTN (Figure S2C) were therefore used to assess mitophagy initiation along with co-staining of Synapsin I (Figures 2D, 2E and S2D). Indeed, mitophagy initiation events marked by mitochondrial EGFP-OPTN foci were predominantly observed within presynapses as indicated by Synapsin I enrichment. To confirm that presynapse mitophagy initiation was driven by the PINK1/Parkin pathway, PINK1 knockout (KO) embryonic stem cell (ESC)-derived iNeurons^15^ (Figure S2E) were analysed. Using the mitophagy initiation marker ATG13, we confirmed that mitophagy within presynapses (as defined by the characteristic bouton morphology (Figure S2F)), was dependent on PINK1 activity (Figures 2F, 2G and S2G). Similar results were observed with OPTN KO neurons (Figures S6A and S6B). These results indicate that PINK1/Parkin mitophagy is the primary responder to oxidative stress induced mitochondrial damage in neurons, despite the presence of high levels of NIX expression (Figure S2B).

**Figure 2.**
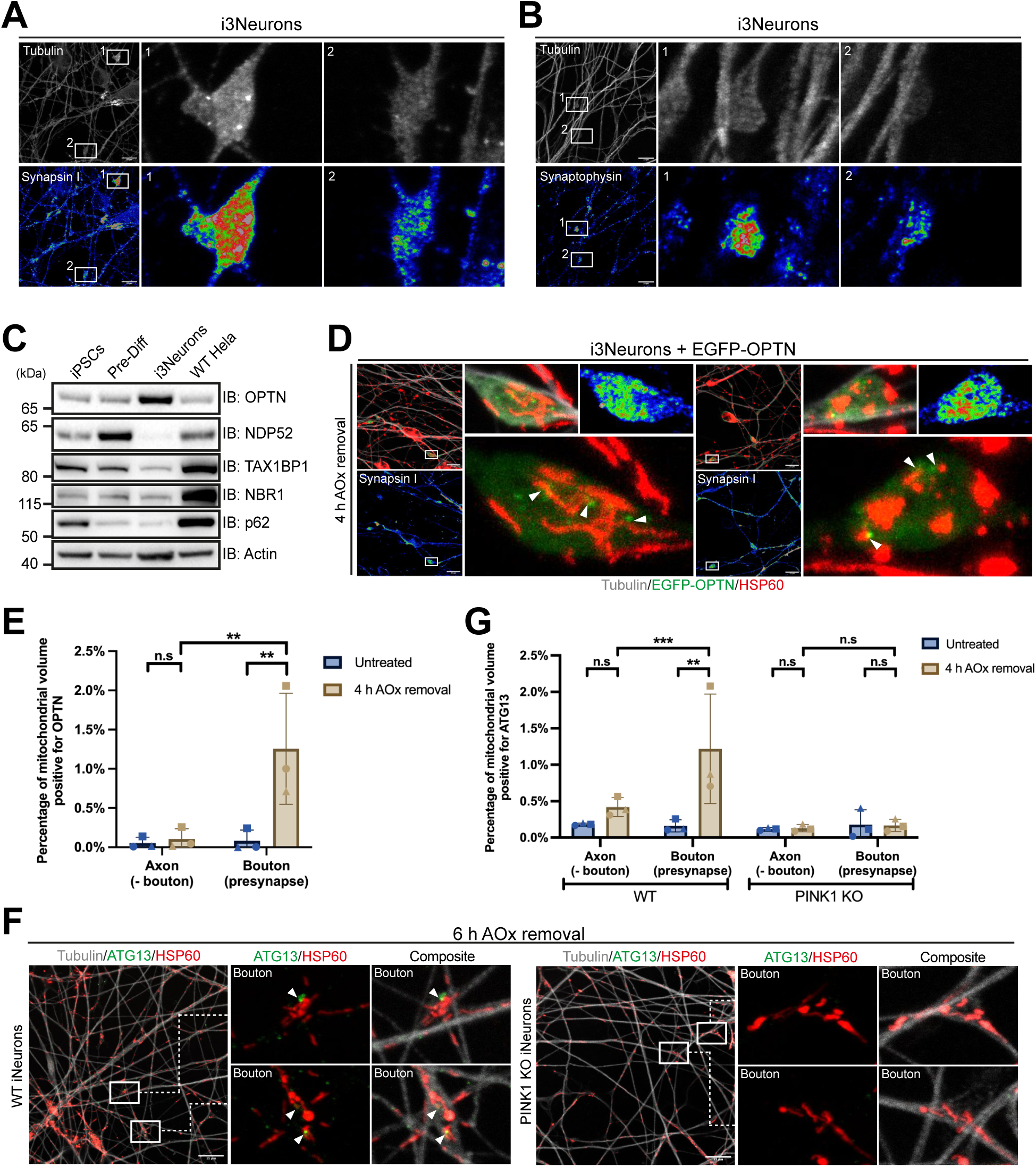
Presynapse is a hotspot for axonal PINK1/Parkin dependent mitophagosome formation (A and B) WT i3Neurons were immunostained for Synapsin I (A) or Synaptophysin (B) and Tubulin. The Synapsin I and Synaptophysin stainings were displayed as RGB thermal spectrum. (C) Immunoblot (IB) for autophagy adaptors including OPTN, NDP52, TAX1BP1, NBR1 and p62 in iPSCs, day 3 pre-differentiated (Pre-Diff) iPSCs, i3Neurons and WT Hela cells. kDa, kilodaltons. (D and E) WT i3Neurons expressing EGFP-OPTN were cultured in AOx removal media for 4 h and immunostained for Synapsin I, EGFP, HSP60 and Tubulin (D) and the percentage of mitochondrial volume positive for OPTN was quantified (E). Untreated samples are shown in Figure S2D. The Synapsin I staining were displayed as RGB thermal spectrum. White arrowheads indicate OPTN puncta localised to mitochondria. (F and G) WT and PINK1 KO iNeurons were cultured in AOx removal media for 6 h and immunostained for ATG13, HSP60 and Tubulin (F) and the percentage of mitochondrial volume positive for ATG13 was quantified (G). Untreated samples shown in Figure S2D. White arrowheads point to ATG13 puncta localised to mitochondria. Data in (E) and (G) are mean ± SD from three independent experiments. Two-way ANOVA. **p<0.005, ***p<0.001. n.s., not significant. Scale bars, 15 µm.

Our i3Neuron and iNeuron differentiations showed enrichment for pre- and post-synaptic markers (Figures 2A, 2B, S2A and S2F), and the NGN2 mediated differentiation approach has been previously shown to generate active glutamatergic excitatory cortical neurons.^43^ However, to further investigate presynaptic mitophagy in a Parkinson’s disease-relevant system, we generated mature iPSC-derived midbrain cultures containing electrophysiologically active neurons using established protocols.^57–59^ To capture individual brain tissue heterogeneity, our cultures included astrocytes and neurons derived from four healthy individuals (7 clones) in the same dish (Figure 3A), consistent with current standards.^60^ We confirmed the midbrain profile of the cultures, with >85% of neurons expressing midbrain markers FOXA2 and LMX1A, and the A9 midbrain profile (GIRK2+, CALB-), relevant to Parkinson’s disease (Figure 3A), consistent with current standard. Approximately 25% of the neurons were dopaminergic (TH+) (Figure 3A), consistent with current standards. To enhance functional and synaptic maturity, neural progenitors were matured in BrainPhys medium on diaminopropane-coated glass coverslips for >90 days before analysis ^57–59^. Patch-clamping analyses confirmed that the neurons exhibited physiological action potentials (Figures 3B and 3C) and received functional excitatory and inhibitory synaptic inputs (Figure 3D). Following confirmation of spontaneous synaptic activity of these neuronal cultures, we assessed presynaptic mitophagy using the autophagy marker WIPI2. In untreated neurons, we observed some evidence of presynaptic mitophagy (Figure 3E, left panel, and S3A), which increased under of oxidative stress conditions (Figure 3E, right panels). These findings indicate that presynaptic mitophagy occurs basally in synaptically active neurons, and that this is intensified under oxidative stress conditions.

**Figure 3.**
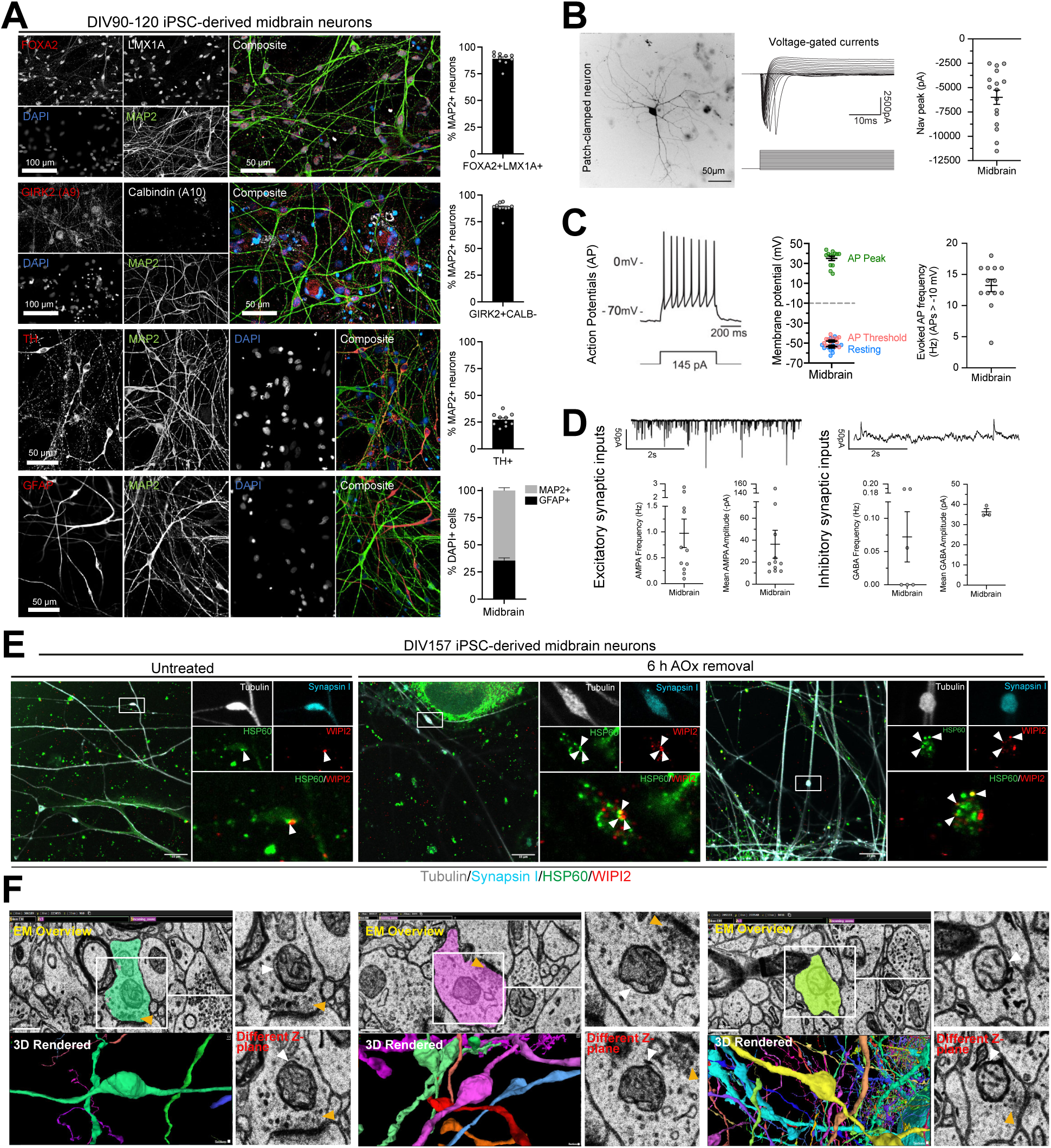
Evidence of presynaptic mitophagy in highly active and mature iPSC-derived midbrain neurons and human brain (A) Representative images (left) and quantification (right) of DIV90-DIV120 human iPSC-derived neuronal cultures expressing dopaminergic (first and third row) and substantia nigra pars compacta (second row) and neurons (MAP2+) versus astrocyte (GFP+; bottom row) identity markers (after 90-120 days in BrainPhys maturation medium. Means ± SEM. Each data point represents one replicate well containing an average of 5600 neurons (MAP2+) across 25 fields of view. (B-D) Whole-cell patch-clamping of healthy human iPSC-derived midbrain neurons after 142-164 days in BrainPhys maturation medium. Mean ± SEM. Each data point represents a single neuron patched. (B) Example of a live human neuron that has been filled with rhodamine from the patch pipette. Midbrain iPSC-derived neurons develop functional voltage-gated sodium channels. (C) Midbrain iPSC-derived neurons develop mature, repetitive action potentials. (D) Midbrain iPSC-derived neurons develop functional excitatory (AMPA) and inhibitory (GABA) postsynaptic circuits. (E) Representative images of DIV157 iPSC-derived midbrain neurons cultured in basal (left) or 6 h of AOx removal condition (middle and right) and immunostained for Synapsin I, WIPI2, HSP60 and Tubulin. White arrowheads indicate WIPI2 puncta localised to mitochondria. (F) Electron microscopy (EM) and the 3D rendered structures images were obtained from the open access dataset generated by Shapson-Coe et al., featuring high-resolution serial section EM of 1 mm^3^ of human temporal cortex ^61^. The presynapses were identified by the distinct bouton morphology along the axon (shown in 3D rendered panel) and the presence of presynaptic density (shown in EM overview or insets, indicated by orange arrowhead). Several examples of mitophagy events were observed in the presynapses, indicated by the autophagosomal membrane forming around the mitochondrion (white arrowhead). Different Z-plane of the EM images (bottom right panel) were examined to identify the completion of the autophagosomal membrane structure.

We next asked whether there is also evidence for presynaptic mitophagy in the human brain. To achieve this, we analysed the rich dataset generated by Shapson-Coe *et al.*,^61^ in which human temporal cortex was imaged by serial section electron microscopy and automated three-dimensional segmentation.^61^ We scanned the open access dataset for presynaptic mitophagy by identifying synaptic sites based on their characteristic morphology of presynaptic vesicle clusters opposed by a postsynaptic density (Figures 3F and S3B, orange arrowheads). Indeed, several examples of presynaptic mitophagy were identified in the human brain, in which a mitochondrion localised within a presynapse had the morphological characteristics of an autophagosomal membrane forming around it (Figures 3F and S3B, white arrowheads). Given that autophagosomal membranes were often not completely sealed around mitochondria (Figures 3F and S3B, see different z-planes), combined with our observations in Figures 1 and 2, we conclude that these are physiological examples of mitophagy initiation within presynapses.

### Anatomy of a presynapse undergoing mitophagy

Autophagosome formation is a complex process that draws on various lipid sources, including ATG9A vesicles and the ER,^6,11,13^ to build and expand autophagosomal membranes around cargoes. We reasoned that axons may lack such machineries, whereas presynaptic sites are rich in lipids and organelles necessary for synaptic transmission, and therefore may be better equipped for building autophagosomes. To determine the membrane architecture of a presynapse undergoing mitophagy, and establish whether a mitophagic presynapse is indeed rich in the membrane sources and ER-phagophore contacts necessary for autophagosome formation, we utilised volumetric electron microscopy. i3Neurons expressing EGFP-OPTN and stained with TMRM, SiR-Tubulin, and Hoechst were subjected to oxidative stress and imaged using live cell microscopy (Figure 4A). EGFP-OPTN was used to identify mitophagy initiation events, while TMRM staining was used to differentiate between healthy or damaged mitochondria. Correlative light and electron microscopy (CLEM) was combined with focused ion beam scanning electron microscopy (FIB-SEM), to locate mitophagy initiation events marked by mitochondrial EGFP-OPTN foci within boutons (Figure 4A), and generate a volumetric electron microscopy dataset (Video S1, upper panel). All bouton membranes were segmented using AIVE (Artificial Intelligence directed Voxel Extraction)^62^ (Figures S4A and S4B, Video S1, lower panel), and used for 3D reconstructions (Figure 4B, Video S2). The 3D reconstruction analysis revealed that the bouton was highly enriched in membranous structures and organelles, including multilamellar structures and ER, demonstrating that it contained the requisite lipid sources for autophagosome formation (Video S2).

**Figure 4:**
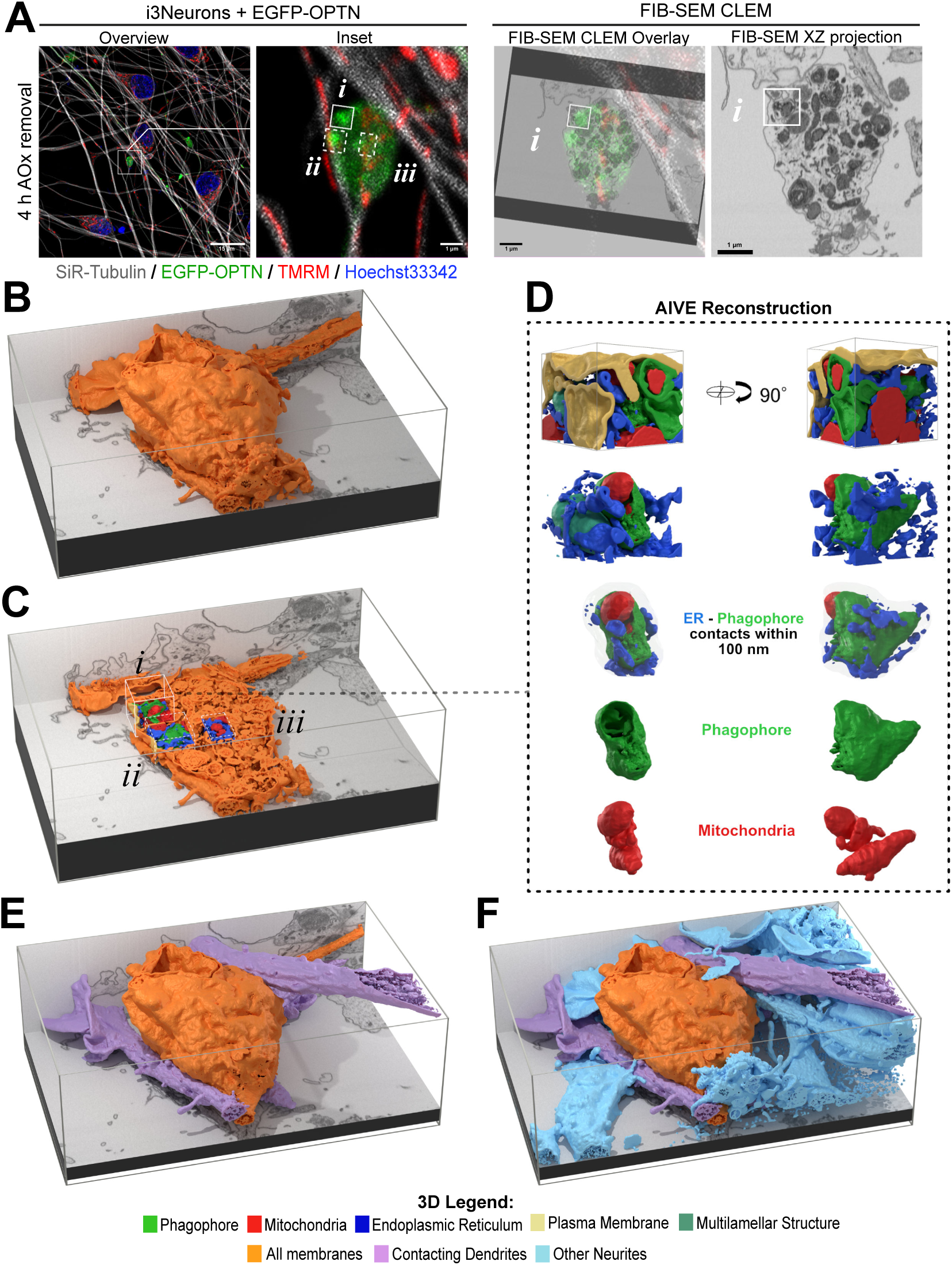
AIVE reconstruction of developing mitophagosome in a presynapse. (A) FIB-SEM CLEM of the identified presynapse with OPTN positive puncta (green). The i3Neuron network as imaged by live confocal microscopy (Overview) and zoomed to the presynapse (Inset). Three areas of interest are identified within this presynapse (*i*, *ii* and *iii*). The confocal and FIB-SEM stacks are then aligned with 3D FIB-SEM CLEM (Overlay) with *i* marked in position. The XZ resliced projection of the FIB-SEM stack is also shown. All membranes in the FIB-SEM image stack were reconstructed using AIVE. (The paired AIVE result with the FIB-SEM image stack were shown in Video S2.) (B) All membranes within the FIB-SEM stack were also reconstructed by AIVE and the surface of the axon bouton structure is displayed (orange). (C) The location of each of the three subset areas are identified in situ within a cut-through of the entire presynapse. (The relating animated 3D rendering of this entire presynapse shown in Video S3.) (D) Inset of the corresponding mitophagosome (*i*) reconstructed in detail with organelles segmented and coloured based on organelle type. This area reveals an almost complete mitophagosome (green) with encapsulated mitochondrial fragments (red). This inset is 1 µm^3^ and is displayed from two angles of view, rotated by 90°. The animated rotation of 3D rendering of mitophagosome (i) was shown in Video S4. Remaining two areas (dashed boxes) are depicted in Figure S5. (E) The presynapse was further segmented into pre- and post-segments, with the main presynapse structure (orange) revealed with three contacting projections (purple), each identified as a dendrite due to the presence of dendritic spines. (F) All remaining structures (cyan) are also shown in position. The relating animated 3D rendering of the presynapse and its surrounding structures were shown in Video S7.

The AIVE reconstructed bouton contained three prominent EGFP-OPTN foci (Figures 4A and 4C, Video S3). Two of the foci lacked TMRM staining (Figure 4A, box i, and Figures S5A, and S5B, box ii), while one retained TMRM staining (Figures S5A, S5B box iii). The EGFP-OPTN foci lacking TMRM staining were revealed to be mature phagophores in the process of completing their encapsulation of damaged mitochondria (Figure 4D and Video S4, and Figure S5C and Video S5). These phagophores were surrounded by extensive ER contacts (Figures 4D and S5C, ER-phagophore contacts panel, and Videos S4 and S5), including contacts at the open mouth of the phagophore, presumably to aid phagophore growth/completion. The phagophore from box i (Video S4) was adjacent to a multilamellar structure, while the phagophore from box ii (Video S5) was nearby a structure morphologically consistent with Golgi membranes, likely trans-golgi network-derived transport carriers.^47^ Both phagophores were also adjacent to additional mitophagy phagophores that we partially segmented (Videos S4 and S5). However, our analyses focused on the phagophores that were aligned most closely with the brightest central point of the OPTN foci (Figure 4A). The third phagophore (Figure S5A and S5B, box iii), was the least mature, and appeared to be a very early intermediate (Figure S5D, Video S6). Like the other phagophores analysed, it too was surrounded by ER (Figure S5D, ER-phagophore contacts panel, Video S6), with a membrane morphology consistent with an ER cradle for the phagophore.^12,13^ Consistent with this being an early mitophagy event, the mitochondrion retained TMRM staining, potentially indicating early mitochondrial damage but not to the point of complete membrane potential loss.

To confirm that the reconstructed bouton, that was rich in mitophagy events, was indeed a presynapse, the surrounding membranes were also reconstructed using AIVE (Figures 4E and 4F, and Video S7). This analysis showed that dendrites, with characteristic morphologies containing dendritic spines, were in direct contact with the bouton (Figure 4E, Video S7), confirming its identity as a presynapse. Surrounding axons/dendrites were also observed (Figure 4F), and we note their relative lack of internal membrane complexity to the presynapse (Videos S1, S2 and S7). Overall, the AIVE 3D electron microscopy analyses demonstrate that presynapses have the requisite membrane sources, including extensive phagophore-ER contacts, and autophagy machineries (Figures 1-4) needed for autophagosome formation. This likely explains why presynapses are the preferred location for mitophagy initiation and autophagosome formation.

### Mitophagy defects result in axonal degeneration

Given that PINK1/Parkin mitophagy activity is targeted toward damaged mitochondria, we asked what the consequence of defective PINK1/Parkin mitophagy might be on axons and presynapses. WT, PINK1 KO and OPTN KO iNeurons from human ESCs,^15^ were subjected to antioxidant removal followed by measurements of mitochondrial reactive oxygen species (ROS) levels using MitoSOX Red (Figures 5A and 5B). WT neurons showed elevated levels of mitochondrial ROS relative to their untreated controls (Figures 5B and S6C). However, neurons lacking PINK1 or OPTN had significantly higher levels of mitochondrial ROS in comparison to WT controls (Figures 5A, 5B and S6C). Given that mitochondria in PINK1 KO and OPTN KO neurons showed significant signs of mitochondrial ROS stress (Figures 5A, 5B and S6C), we investigated what the effects of prolonged oxidative stress might be. Severely damaged mitochondria undergo mitochondrial outer membrane permeabilization (MOMP) that results in cytochrome *c* release and mitochondrial DNA (mtDNA) release. Cytochrome *c* release activates downstream caspases including caspase 3 which degrade cells during apoptosis,^63–66^ whereas mtDNA release has been linked to inflammation via the cGAS-STING pathway.^67^

**Figure 5.**
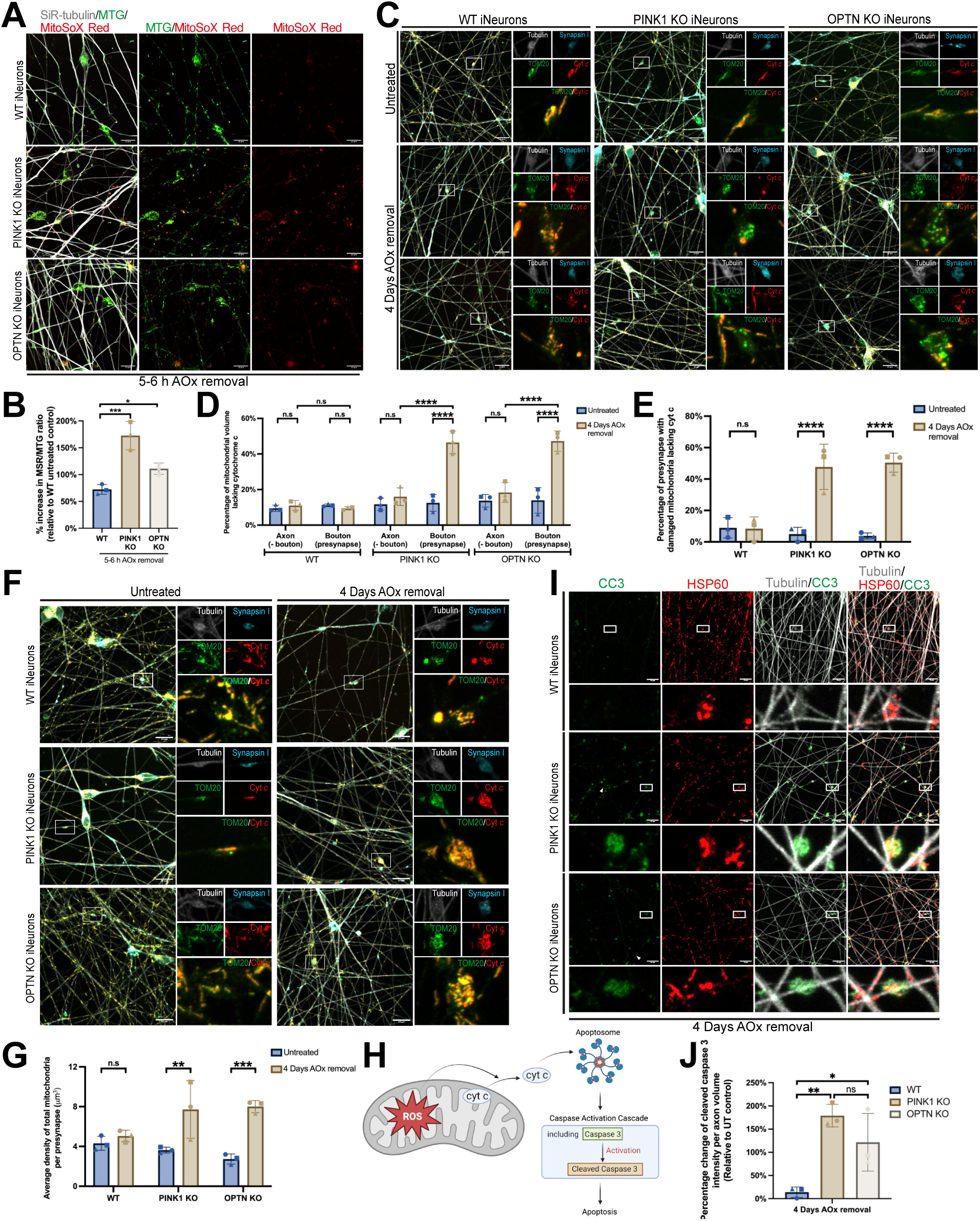
PINK1/Parkin mitophagy protects neurons from oxidative stress-induced mitochondrial collapse and neuronal degeneration (A and B) WT, PINK1 KO and OPTN KO iNeurons were cultured in AOx removal media for 5-6 h and stained with MitoTracker Green (MTG), MitoSOX red (MSR) and SiR-Tubulin (A) and the percentage change in MSR to MTG ratio compared to untreated control was quantified (B). Untreated samples are shown in Figure S6C. (C-E, F-G) WT, PINK1 KO and OPTN KO iNeurons were treated with pan-caspase inhibitor QVD-OPh (QVD) and cultured in untreated or AOx removal conditions for 4 days, and were immunostained for Synapsin I, TOM20, Cytochrome *c* (Cyt *c*) and Tubulin (C, F). The percentage of mitochondrial volume lacking Cyt *c* (D), the percentage of the number of presynapse with damaged mitochondria lacking Cyt *c* (E) and the average density of total mitochondria per presynapse (G) were quantified. (H) Schematic of the mechanism of reactive oxygen species (ROS) activating apoptosis. (I and J) WT, PINK1 KO and OPTN KO iNeurons were cultured in AOx removal media for 4 days and were immunostained for Cleaved Caspase 3 (CC3), HSP60 and Tubulin (I). The fold change of CC3 intensity per axon volume (J) was quantified. Untreated samples are shown in Figure S6D. White arrowheads point to axons with caspase 3 activation. Data in (B), (D), (E), (G) and (J) are mean ± SD from three independent experiments. Two-way ANOVA. p*<0.05, **p<0.005, ***p<0.001, ****p<0.0001. n.s., not significant. Scale bars, 15 µm.

To assess MOMP events, WT, PINK1 KO, and OPTN KO neurons were subjected to AOx removal treatment for 4 days. This treatment was combined with the pan-caspase inhibitor QVD-OPh (Quinoline-Val-Asp-Difluorophenoxymethylketone) to capture MOMP events that can be lost following caspase-mediated proteolysis. First, we analysed cytochrome *c* release in presynapses by co-staining neurons for the mitochondrial protein TOM20 and cytochrome *c* (Figures 5C and 5D). Mitochondria undergoing cytochrome *c* release will remain positive for TOM20 but will become negative for cytochrome *c*. WT controls did not show evidence of oxidative stress induced cytochrome *c* release following 4 days of antioxidant removal (Figures 5C, 5D and 5E). In contrast, PINK1 KO and OPTN KO neurons showed significantly increased levels of cytochrome *c* release, specifically within presynapses (Figures 5C, 5D and 5E). Overall, approximately 50% of PINK1 KO and OPTN KO presynapses had evidence of MOMP (Figure 5E). Notably, no significant increase in cytochrome *c* release was observed in axonal regions outside of presynapses in PINK1 KO, or OPTN KO iNeurons (Figures 5C and 5D).

During the cytochrome *c* release analyses in Figures 5C-E, we noted that PINK1 KO and OPTN KO presynapses lacking MOMP appeared to have increased mitochondrial density. Given that PINK1/Parkin activation can occur along the length of axons, but mitophagy predominantly occurs in presynapses (Figure 1 and 2), it is likely that damaged axonal mitochondria are transported to presynapses to undergo mitophagy. Neurons defective in PINK1/Parkin mitophagy may therefore accumulate mitochondria within presynapses. To address this point, we analysed the ∼50% of presynapses from Figure 5C-E, that did not yet show evidence of MOMP following 4 days of oxidative stress. Indeed, in these presynapses, we observed a significant accumulation of mitochondria in PINK1 KO and OPTN KO neurons, but not in WT controls (Figures 5F and 5G), consistent with damaged axonal mitochondria being transported to presynapses for mitophagy.

Given that cytochrome *c* is released locally within presynapses of PINK1/Parkin mitophagy defective neurons, we assessed whether there was also evidence for downstream caspase-3 activation (Figure 5H). WT, PINK1 KO, and OPTN KO iNeurons were subjected to 4 days of AOx removal this time in the absence of caspase inhibitor and stained with a cleaved caspase-3 specific antibody (Figures 5I and S6D). PINK1 KO and OPTN KO neurons showed a substantial increase in cleaved caspase 3 staining (Figures 5I and 5J), predominantly in regions with bouton morphology, although caspase 3 activation was also observed along the length of some axons (Figure 5I, white arrowheads). In contrast, the WT control showed little to no caspase activation, consistent with their lack of cytochrome c release (Figures 5I, 5J and S6D), demonstrating that PINK1/Parkin mitophagy is neuroprotective against oxidatively damaged mitochondria. Collectively, these results show that defects in mitophagy result in damaged mitochondria accumulating within presynapses, where they locally release cytochrome c and activate degradative caspases.

MOMP events resulting in herniation of the mitochondrial inner membrane have been shown to release mtDNA.^68,69^ We asked whether mtDNA might also be released along with cytochrome *c* in mitophagy defective neurons, potentially resulting in downstream inflammatory signalling.^67,70^ However, despite observing evidence for mtDNA release using the positive control treatment of BH3 mimetic ABT-737 ^68^ (Figure S7A), we did not observe evidence for mtDNA release in PINK1 KO or OPTN KO neurons following 4 days of oxidative stress (Figure S7A). In addition, there was little to no expression of STING observed in the iNeurons (Figure S7B). These results indicate that while oxidative stress induced MOMP is sufficient to release cytochrome *c*, it does not promote mtDNA release through inner membrane herniation, at least not under the conditions tested.

### HIF1α-mediated mitophagy through prolyl-4 hydroxylase inhibition rescues axon degeneration in PINK1/Parkin defective neurons

Given that alternative forms of mitophagy were not strongly stimulated in response to oxidatively damaged mitochondria in PINK1 KOs (Figures 2F and 2G), we asked whether synthetically stimulating alternative mitophagy might be beneficial in PINK1/Parkin defective neurons. Mitophagy receptors such as BNIP3 and NIX drive an alternative form of mitophagy that can be stimulated by HIF1α signalling,^27,71^ which is negatively regulated by prolyl hydroxylases.^72,73^ Small molecule prolyl-4 hydroxylase inhibitors have been used in the clinic to activate HIF1α signaling,^74,75^ and one of these, Roxadustat, was tested for its ability to drive mitophagy in neurons. PINK1 KO, OPTN KO, and WT controls were treated either with Roxadustat for 48 h, AOx removal for 4 days, or a combination of antioxidant removal for 4 days with Roxadustat treatment in the final 48 h (Figures 6A, 6B, S8A and S8C). Roxadustat treatment successfully induced mitophagy not only in WT controls, but also in PINK1 KO and OPTN KO neurons (Figure 6A and 6B). Combination treatment of oxidative stress and Roxadustat resulted in much higher mitophagy levels in PINK1 KO and OPTN KO neurons (Figures 6A, 6B and S8A), perhaps due to higher levels of damaged mitochondria in the KO lines, or due to increased ROS-mediated promotion of HIF1α signaling.^76^

**Figure 6.**
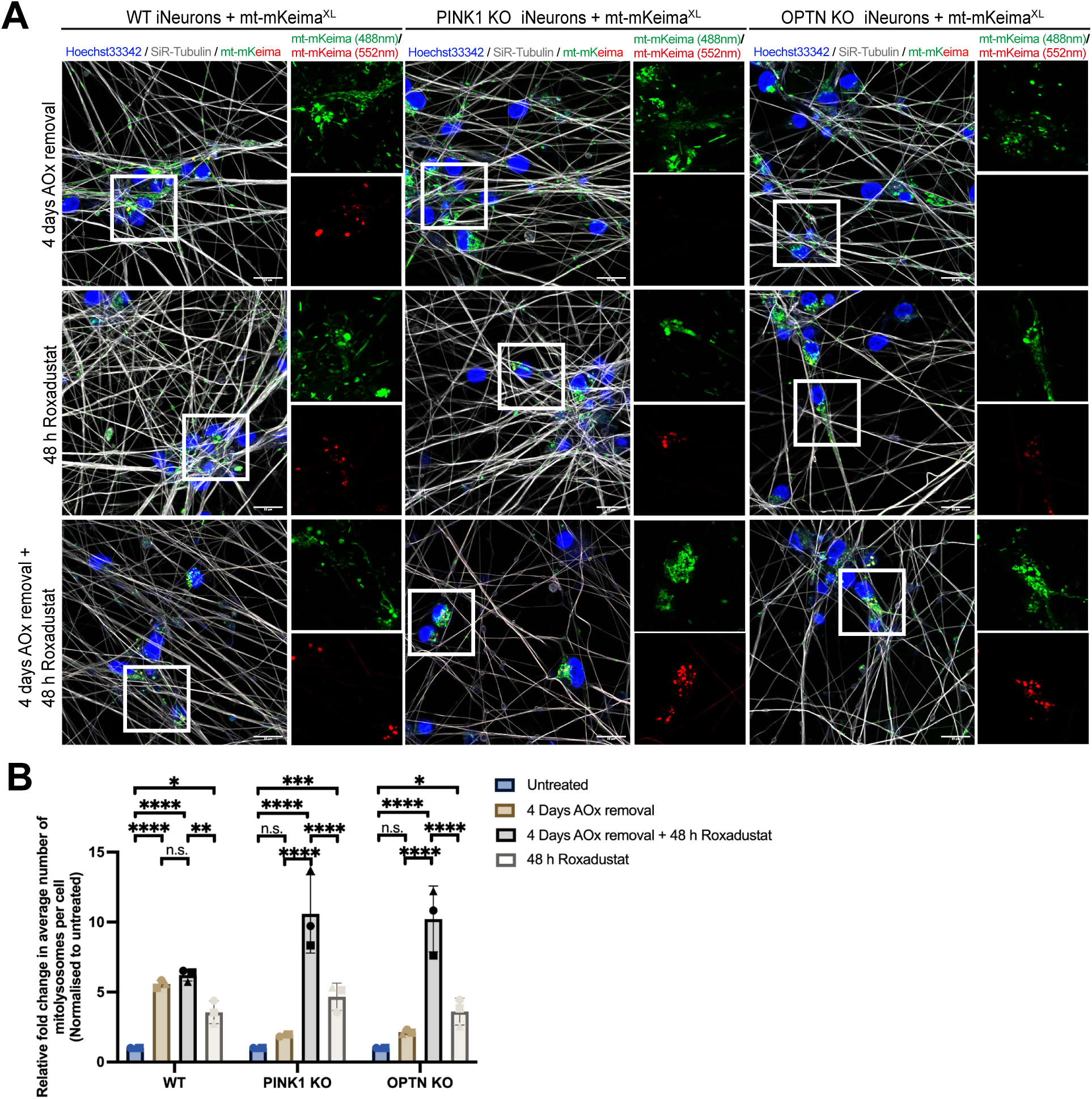
Roxadustat stimulates PINK1-independent mitophagy in PINK1/Parkin-mitophagy defective neurons in response to oxidatively damaged mitochondria. (A and B) WT, PINK1 KO and OPTN KO iNeurons expressing mito-mKeima^XL^ (mt-mKeima^XL^) were treated with pan-caspase inhibitor QVD-OPh (QVD) and cultured in either AOx removal media for 4 days, 40 µM of Roxadustat for 48 h, or a combination of AOx removal for 4 days with Roxadustat treatment in the final 48 h. These neurons were stained with SiR-Tubulin and Hoechst33342 and underwent live cell imaging. Untreated samples are shown in Figure S8A. The relative fold change in average number of mitolysosomes per cell were quantified in (B). Data in (B) is mean ± SD from three independent experiments. Two-way ANOVA. *p<0.05, **p<0.005, ***p<0.001, ****p<0.0001. n.s., not significant. Scale bars, 15 µm.

We next determined whether mitophagy induced by Roxadustat could restore mitochondrial health and prevent axon degeneration. Mitochondria were stressed for 4 days with antioxidant removal and were either treated with or without Roxadustat in the final 48 h (Figures 7 A-D and S8B). WT neurons did not show evidence of caspase-3 activation, neither with 4 days of antioxidant removal nor with Roxadustat treatment, arguing that Roxadustat is not toxic to neurons. In contrast, PINK1 KO and OPTN KO neurons showed significant levels of caspase-3 activation via imaging (Figures 7A, 7B and S8B) and western blot analysis (Figure 7C and 7D). In addition, PINK1 KO and OPTN KO neurons showed evidence of synapse loss based on the reduction of both Synapsin I and PSD-95 (Figure 7C). Strikingly, Roxadustat treatment almost completely reversed caspase 3 activation and synapse loss (Figures 7C and 7D). Consistently, Roxadustat also prevented presynaptic cytochrome c release (Figures S9A, S9B and S9C) and mitochondrial accumulation within presynapses (Figures S9D and S9E). This result argues that presynapses are universal mitophagy sites that are not restricted only to PINK1/Parkin mitophagy. Overall, these results demonstrate that stimulation of alternative mitophagy via HIF1α prolyl-4 hydroxylase inhibition can clear damaged mitochondria and protect axons from oxidative stress, revealing a potential therapeutic avenue. In neurons, BNIP3 may be the primary driver given its strong response to Roxadustat (Figure 7C).

**Figure 7.**
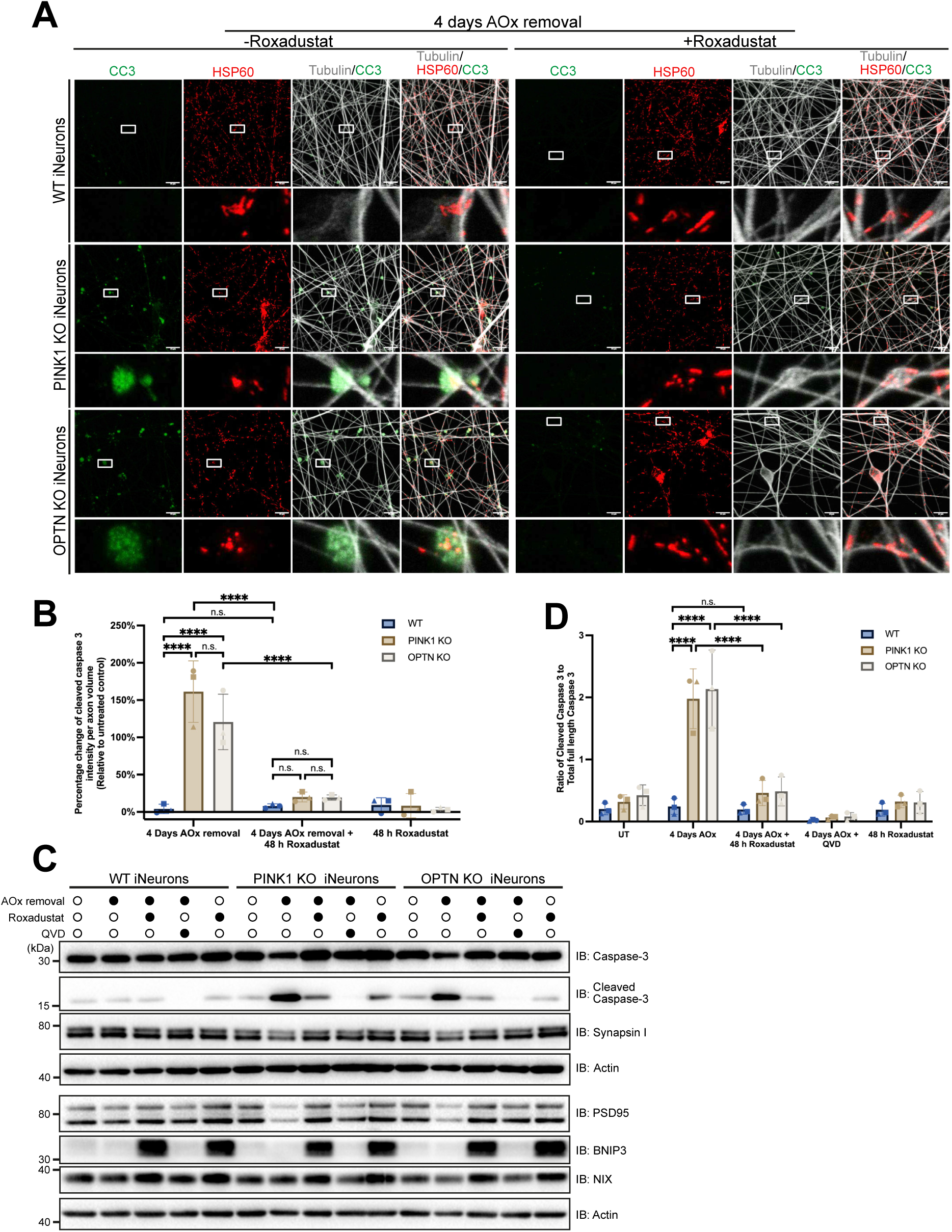
Roxadustat restores mitochondrial health in PINK1/Parkin-mitophagy defective neurons and prevents axon degeneration. (A and B) WT, PINK1 KO and OPTN KO iNeurons were treated with either AOx removal media for 4 days or a combination of AOx removal for 4 days with 40 µM Roxadustat treatment in the final 48 h. The neurons were immunostained for Cleaved Caspase 3 (CC3), HSP60 and Tubulin (A). The percentage change of CC3 intensity per axon volume (B) was quantified. Untreated and 48 h 40 µM Roxadustat treated samples shown in Figure S8B. (C and D) The activation of Caspase-3, levels of synaptic proteins Synapsin I and PSD95, and the expression of PINK1/Parkin-independent mitophagy adaptor proteins BNIP3 and NIX were analysed through immunoblotting in WT, PINK1 KO and OPTN KO iNeurons that were untreated, treated with either AOx removal condition for 4 days, 40 µM Roxadustat for 48 h, a combination of AOx removal for 4 days with 40 µM Roxadustat treatment in the final 48 h, or QVD. The ratio of Cleaved Caspase-3 to total full length Caspase-3 were quantified (D). Data in (B) and (D) were mean ± SD from three independent experiments. Two-way ANOVA. ***p<0.001, ****p<0.0001. n.s., not significant. Scale bars, 15 µm.

## Discussion

Mitochondria play essential roles in neuronal homeostasis, but the polarized architecture of neurons in which mitochondria can become damaged in a distal location creates challenges for mitochondrial quality control. Mitophagy mediated by PINK1/Parkin has been reported in murine axons,^42,77^ but its axonal importance and spatial requirements, if any, remained open questions.

By exploring the cell biology of mitophagy in human neurons, with a focus on axons, we uncovered a role for presynapses as mitophagy pit stops (Figure S10A). Given that HIF1α stimulated alternative mitophagy and cleared damaged mitochondria within presynapses of PINK1 and OPTN KOs (Figures 6 and 7), we conclude that presynaptic pit stop mitophagy is a universal mechanism that is not restricted only to the PINK1/Parkin pathway. Our data are consistent with the following model (Figure S10A): Distally damaged axonal mitochondria, including those labelled by PINK1/Parkin, are transported along axons, and upon encountering an *en passant* presynapse, mitochondrial transport is stalled so that an autophagosome may be built around the damaged mitochondrion (Figure S10A). While the mechanism of trapping damaged mitochondria within presynapses for mitophagy is unclear, factors that anchor mitochondria in axons including syntaphilin,^78^ the LKB1-NUAK1 kinase pathway,^79^ or modulation of mitochondrial trafficking machineries such as Miro^80,81^ may play a role. Presynapses are highly permissive sites for axonal mitophagy because they contain the autophagy machineries, including extensive lipid sources, necessary for building autophagosomes (Figures 1, 4 and S10). This is consistent with prior studies reporting enrichment of ATG9A in presynaptic terminals, in which autophagy activity has been reported. ^8,82^ Notably, mitochondrial degradation does not occur within presynapses since acidified mitolysosomes were only observed in the soma or axon hillock (Figures 1B, 6A and 6B), in agreement with previous work showing retrograde transport is required for lysosome maturation.^46^ However, since mitochondria are now packaged within the delimiting membrane of an autophagosome/mitolysosome, the risk of MOMP is no longer a danger to neurons. Presynaptic mitophagy therefore provides a critical pathway for distal mitochondrial quality control that avoids the need to transport damaged mitochondria long distances toward the soma for mitophagy.

Presynaptic mitophagy has benefits for mitochondrial quality control in distal axons, but it also creates a point of vulnerability when mitophagy is defective (Figure 5). The accumulation of damaged mitochondria within presynapses that release cytochrome *c* and cause caspase activation represents a potential mechanism for neurodegeneration in disease (Figure S10B). Indeed, it is strikingly consistent with the die-back pathology that has been reported in neurodegenerative diseases including Parkinson’s disease and Alzheimer’s disease, in which axons and their synapses degenerate before the soma.^33–36^ This is typically associated with caspase activation including caspase 3.^83,84^ It is also interesting to note that synaptic caspase 3 activation also occurs in non-pathological settings,^85–87^ where the selective pruning of synapses and axons promotes development and maintains plasticity. However, the existence of these degradative caspase machineries within axons and synapses for physiological purposes can also become pathologically activated when mitophagy is defective.

In neurons lacking PINK1 or OPTN, we observed little to no mitophagy activity during oxidative stress (Figures 2F, 2G, 6A, and 6B), demonstrating that the PINK1/Parkin pathway is the primary responder to damaged mitochondria. This created a therapeutic opportunity to re-activate mitophagy through alternative pathways. In this study we demonstrated one such therapeutic avenue via HIF1α prolyl-4 hydroxylase inhibition using the clinically approved small molecule Roxadustat. Other compounds in this class, including Daprodustat which is FDA approved for anemia in chronic kidney disease,^88^ also hold potential as a mitophagy therapeutic. However, we anticipate that stimulation of HIF1α mediated mitophagy through prolyl-4 hydroxylase inhibition will have benefits beyond conditions in which PINK1/Parkin mitophagy is defective. For example, promoting mitophagy through small molecule mediated USP30 inhibition limited a-synuclein induced neuronal toxicity in murine models.^89^ Moving forward, we hypothesise that boosting mitophagy through Roxadustat (or other prolyl-4 hydroxyl inhibitors such as Daprodustat) could have broad benefits for any disease in which clearance of damaged mitochondria might be beneficial.

---

**Figure S1.**
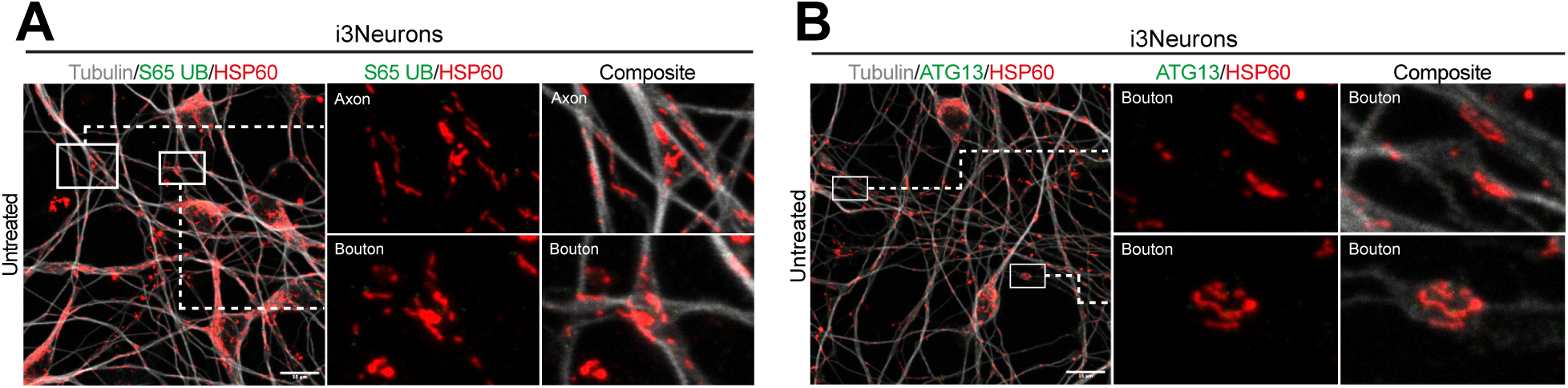
Antioxidant removal damages mitochondria and promotes mitophagy in iPSC-derived i3Neurons (A) Representative confocal images of untreated WT i3Neurons immunostained for S65 UB, HSP60 and Tubulin. Related to Figure 1C and 1D. (B) Representative confocal images of untreated WT i3Neurons immunostained for ATG13, HSP60 and Tubulin. Related to Figure 1E and 1F. Scale bars, 15 µm.

**Figure S2.**
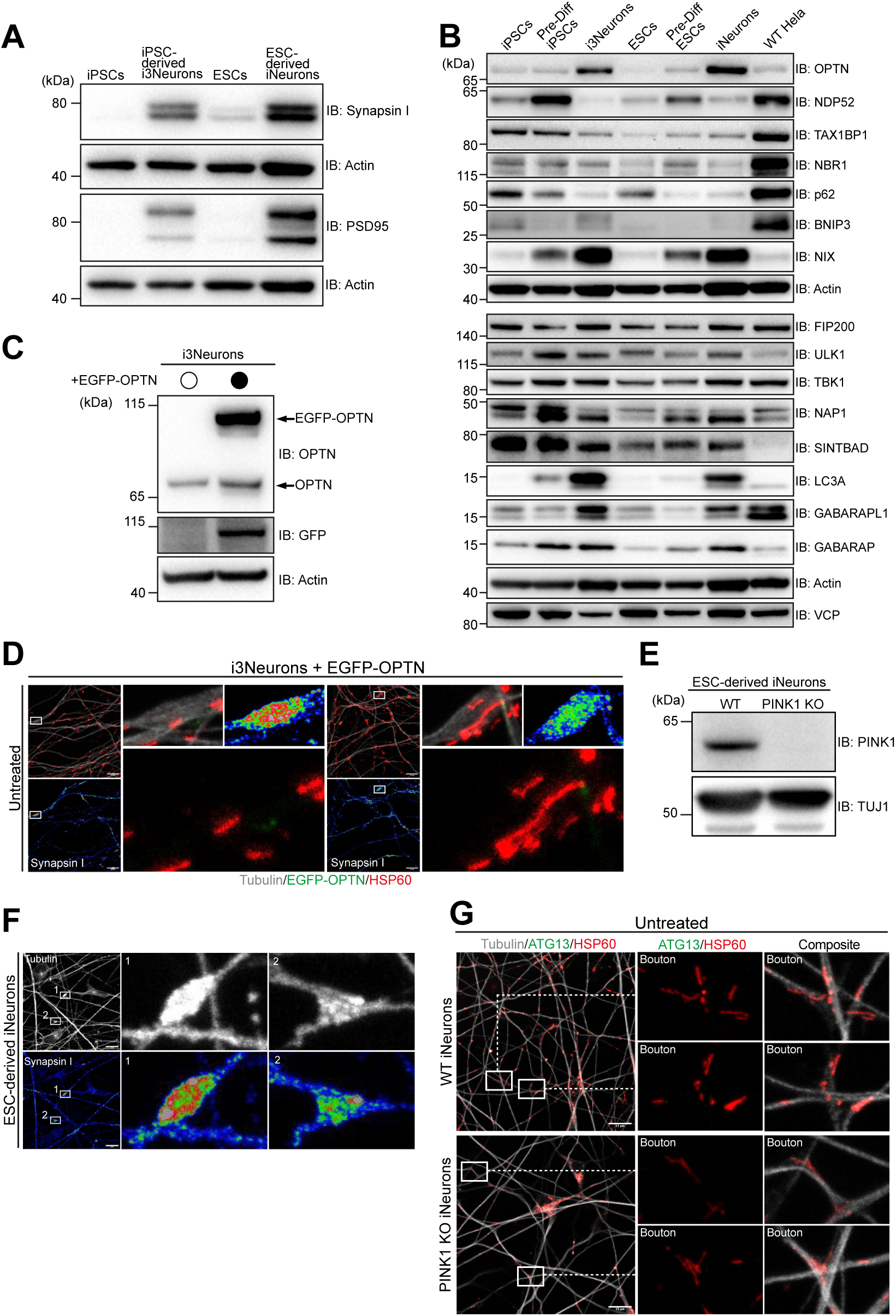
The initiation of PINK1/Parkin dependent mitophagosome formation primarily occurs in presynapses (A) The expression of presynaptic marker Synapsin I and postsynaptic marker PSD95 were examined in iPSCs, iPSC-derived i3Neurons, ESCs and ESC-derived iNeurons using immunoblotting (IB). (B) Immunoblot for indicated autophagy adaptors and regulators in iPSCs, day 3 pre-differentiated (Pre-Diff) iPSCs, iPSC-derived i3Neurons, ESCs, day 7 Pre-Diff ESCs, ESC-derived iNeurons and WT Hela cells. Second from the bottom actin panel is the loading control for p62, FIP200, ULK1, TBK1 and GABARAPL1. The VCP panel is the loading control for NAP1, SINTBAD and LC3A. kDa, kilodaltons. Related to Figure 2C. (C) The expression of EGFP-OPTN in WT i3Neurons was confirmed by IB for OPTN and EGFP. (D) Representative images of untreated WT i3Neurons expressing EGFP-OPTN that were immunostained for Synapsin I, EGFP, HSP60 and Tubulin. The Synapsin I staining is displayed as RGB thermal spectrum. (E) Confirmation of PINK1 KO ESC-derived iNeurons by IB. (F) WT ESC-derived iNeurons were immunostained for Synapsin I. (G) Untreated WT and PINK1 KO iNeurons were immunostained for ATG13, HSP60 and Tubulin. Related to Figure 2F and 2G. Scale bars, 15 µm.

**Figure S3.**
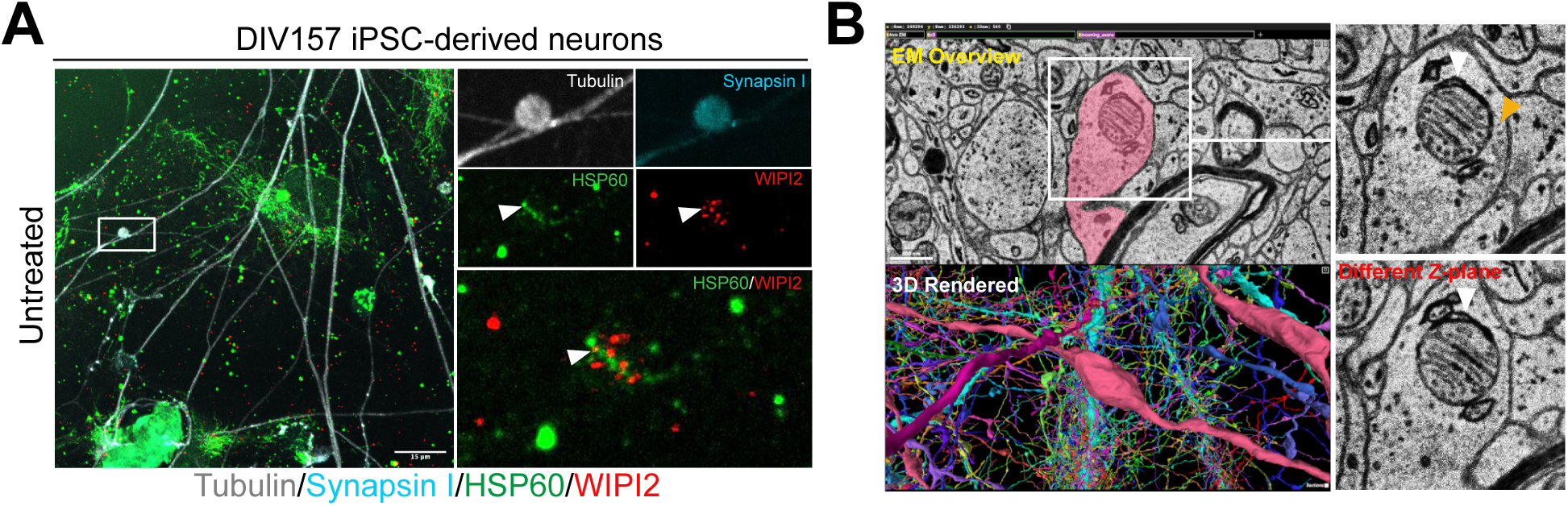
Evidence of presynaptic mitophagy in highly active and mature iPSC-derived midbrain neurons and human brain (A) Representative images of DIV157 midbrain iPSC-derived neurons cultured in basal condition and immunostained for Synapsin I, WIPI2, HSP60 and Tubulin. White arrowhead indicates WIPI2 puncta localised to mitochondria. Related to Figure 3E. (B) Another example electron microscopy (EM) images of presynaptic mitophagy shown in the open access dataset generated by Shapson-Coe et al., featuring high-resolution serial section EM of 1 mm^3^ of human temporal cortex. ^61^ The presynapses were identified by the distinct bouton morphology along axon (shown in 3D rendered panel) and the presence of presynaptic density (shown in EM overview or insets, indicated by orange arrowhead). Mitophagy was indicated by the autophagosomal membrane forming around the mitochondrion (white arrowhead). Different Z-plane of the EM images were examined to identify the completion of the autophagosomal membrane structure. Related to Figure 3F.

**Figure S4.**
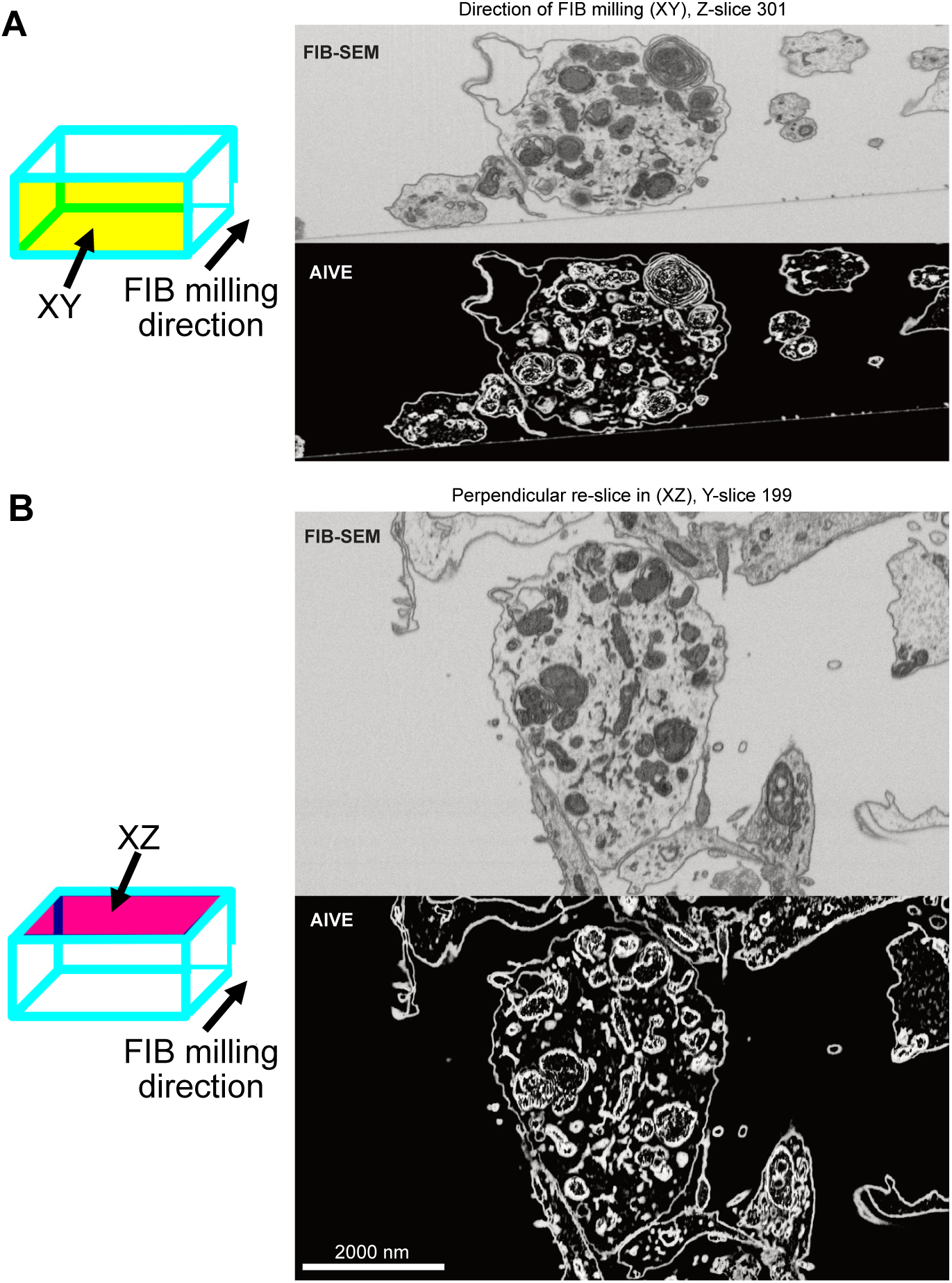
AIVE results paired to the corresponding FIB-SEM image slice. (A) The 3D FIB-SEM image stack was imaged with axes XY with the FIB-SEM (top) and raw AIVE output (bottom) shown for a single slice in the XY direction. This is the same as the FIB milling direction. (The progressive cut-through of the FIB-SEM slices paired with the corresponding AIVE results were shown in Video S1). (B) The XZ resliced projection of the FIB-SEM and raw AIVE output for a single slice of the image stack. This direction corresponds to the Confocal plane but maintains the raw FIB-SEM voxels orientation.

**Figure S5.**
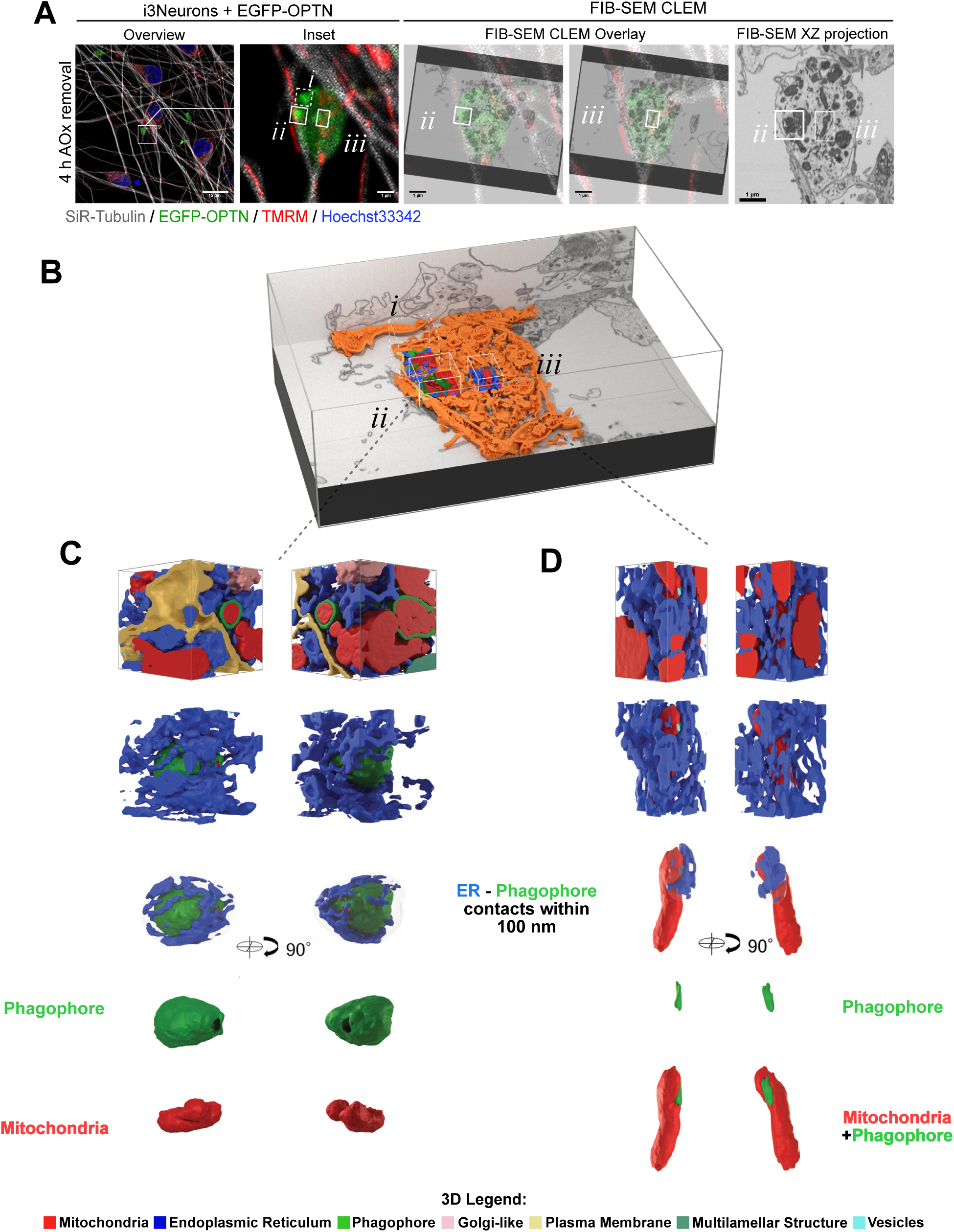
Additional areas of AIVE reconstruction (A) FIB-SEM CLEM of the i3Neuron network with image overview, zoomed in view of axon bouton (Inset), with the three areas (*i*, *ii* and *iii*) marked. FIB-SEM CLEM overlay alignment of *ii* and *iii* with the position of each marked by the solid boxes. An XZ FIB-SEM slice is also shown to locate the two areas. (B) The two areas are displayed in situ within the AIVE reconstruction of the whole presynapse. The two areas include (C) An almost complete mitophagosome, and (D) a mitochondrion with a newly developing phagophore. Each is displayed from two viewing angles separated by 90° rotation.

**Figure S6.**
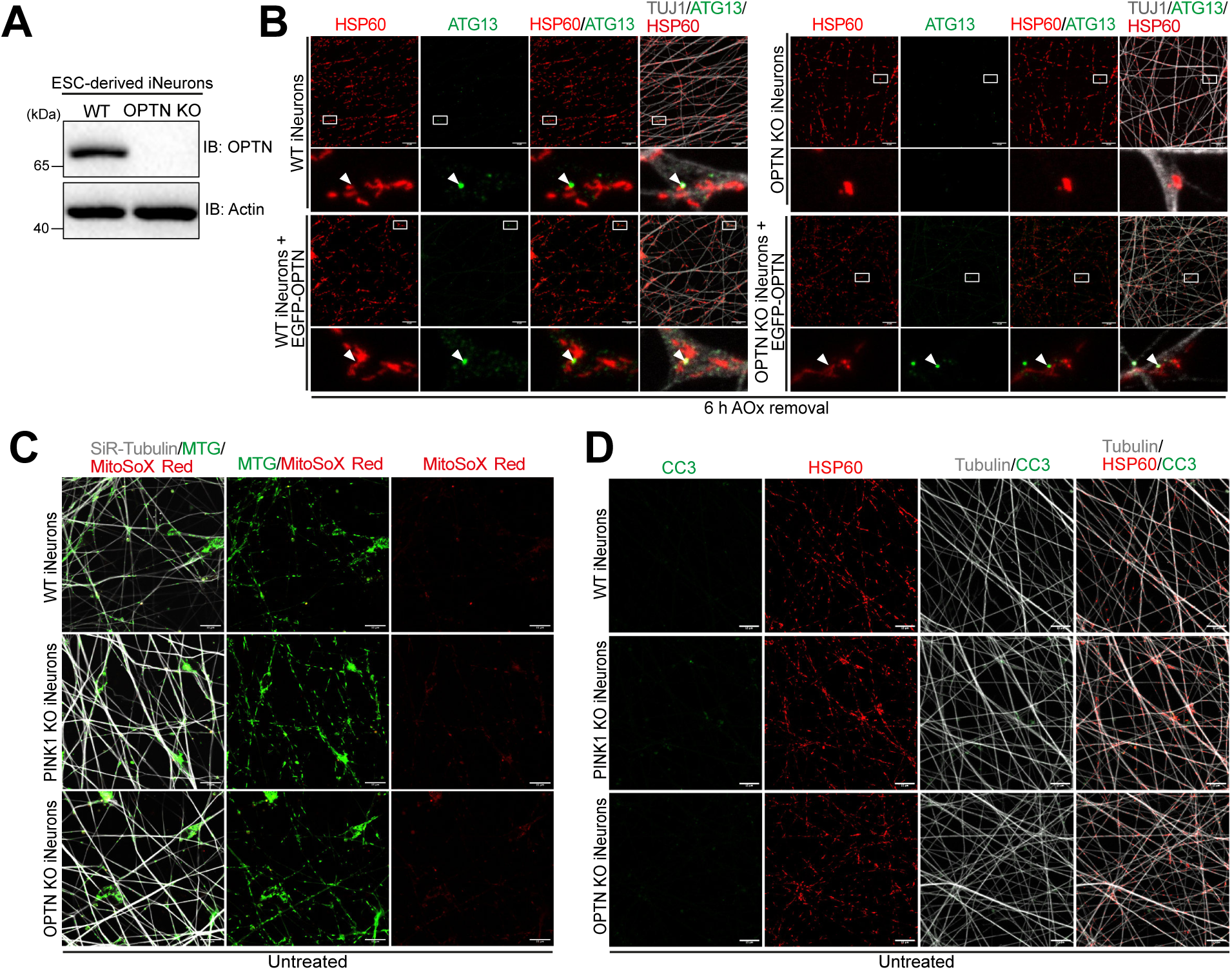
Confirmation that AOx removal-induced PINK1/Parkin mitophagy is OPTN dependent, and PINK1/Parkin mitophagy prevents axon degeneration. (A) Confirmation of OPTN KO ESC-derived iNeurons by immunoblotting. (B) Representative confocal images of WT and OPTN KO iNeurons with or without EGFP-OPTN expression cultured in AOx removal media for 6 h and immunostained for ATG13, HSP60 and TUJ1. White arrowheads point to ATG13 puncta localised to mitochondria. (C) Representative confocal images of untreated WT, PINK1 KO and OPTN KO iNeurons stained with MitoTracker Green (MTG), MitoSOX red (MSR) and SiR-Tubulin. Related to Figure 5A and 5B. (D) Representative confocal images of untreated WT, PINK1 KO and OPTN KO iNeurons immunostained for Cleaved Caspase 3 (CC3), HSP60 and Tubulin. Related to Figure 5I and J. Scale bars, 15 µm.

**Figure S7.**
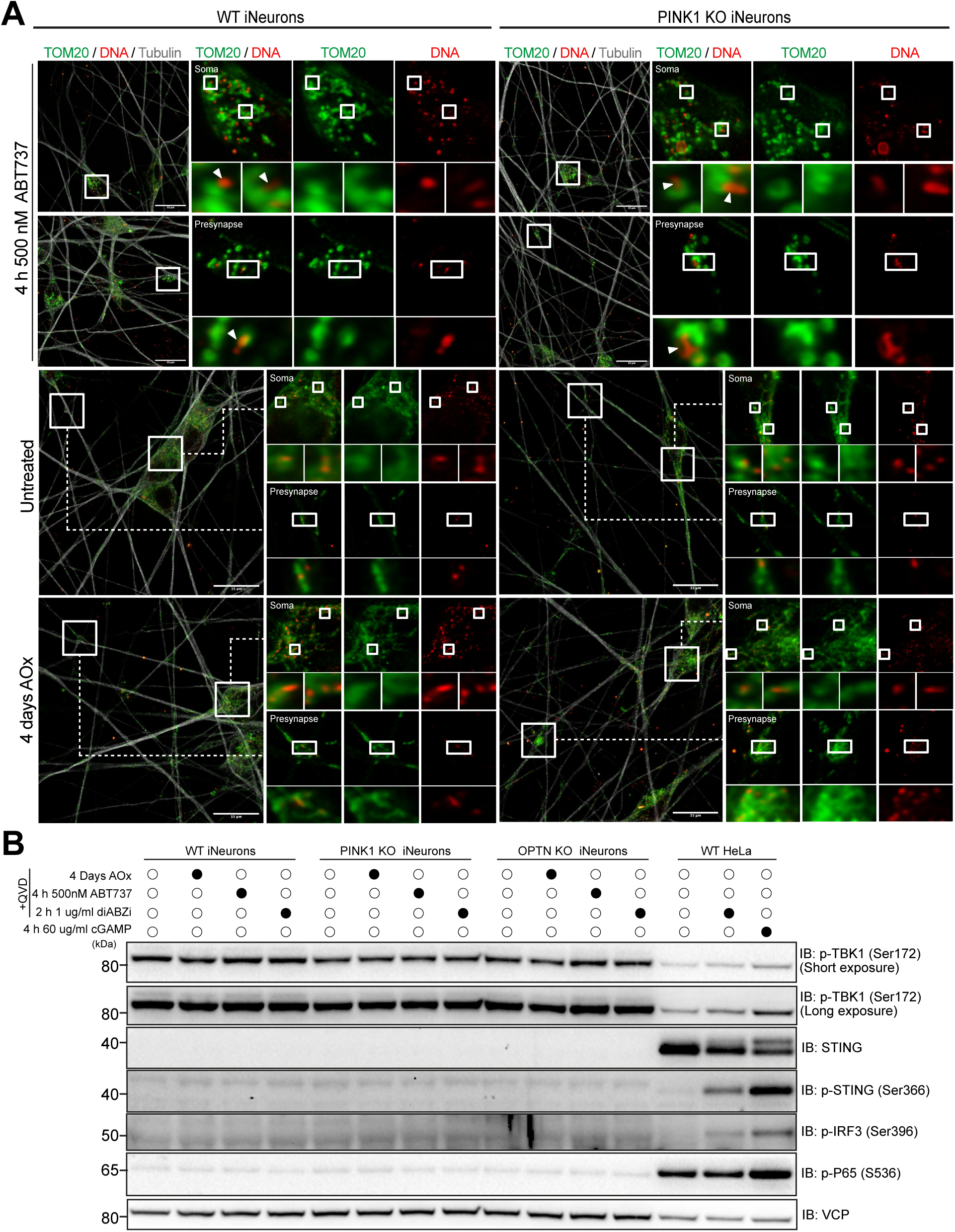
Long term AOx removal does not induce mitochondria herniation or cGAS-STING activation. (A) Representative confocal images of WT and PINK1 KO iNeurons that were untreated, treated with 500 nM ABT737 for 4 h or AOx removal for 4 days and immunostained for TOM20, DNA and Tubulin. White arrowheads point to herniated mitochondria with mtDNA leakage. The overview images were captured using 40X objective with 2.5X zoom and the inset images were captured separately using the same objective with 6.5X zoom. Scale bars, 15 µm. (B) WT, PINK1 KO, OPTN KO iNeurons and WT HeLa cells were untreated, treated with AOx removal for 4 days, 500 nM ABT737 for 4 h, 1 µg/ml diABZi for 2 h or 60 µg/ml cGAMP for 4 h, and immunoblotted for regulators of cGAS-STING pathway including p-TBK1 (Ser172), STING, p-STING (Ser366), p-IRF3 (Ser396), p-P65 (S536).

**Figure S8.**
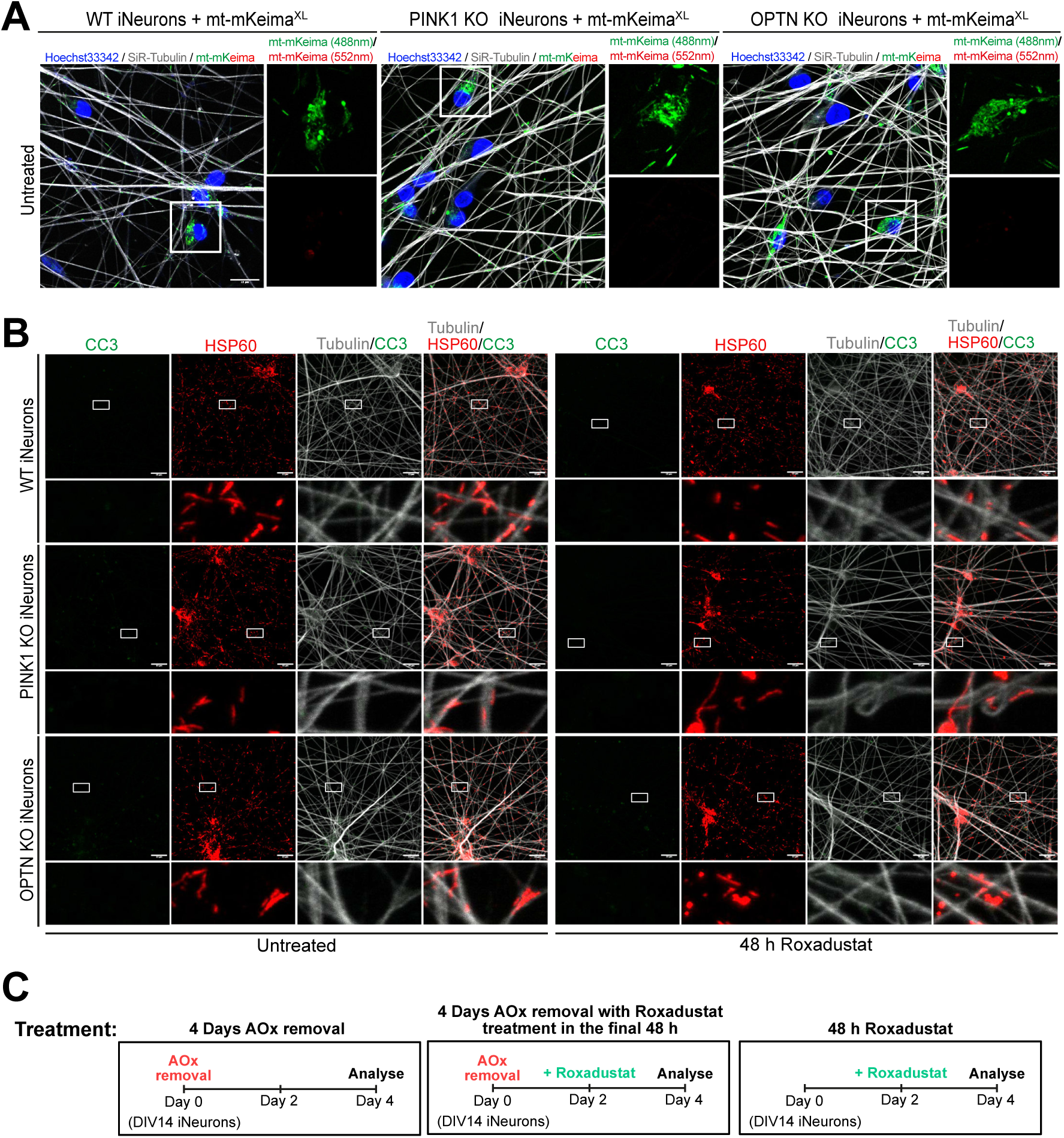
Roxadustat protects PINK1/Parkin-mitophagy defective neurons from axon degeneration by restoring mitochondrial health. (A) Representative live cell confocal images of QVD treated control WT, PINK1 KO and OPTN KO iNeurons expressing mt-mKeima^XL^ stained with SiR-Tubulin and Hoechst33342. Related to Figure 6A and 6B. (B) Representative confocal images of untreated and 48 h 40 µM Roxadustat treated WT, PINK1 KO and OPTN KO iNeurons immunostained for Cleaved Caspase 3 (CC3), HSP60 and Tubulin. Related to Figure 7A and 7B Scale bars, 15 µm. (C) Experimental overview of how the 4 days AOx removal, a combination of 4 days AOx removal with 40 µM Roxadustat treatment in the final 48 h and the 48 h 40 µM Roxadustat treatment were performed.

**Figure S9.**
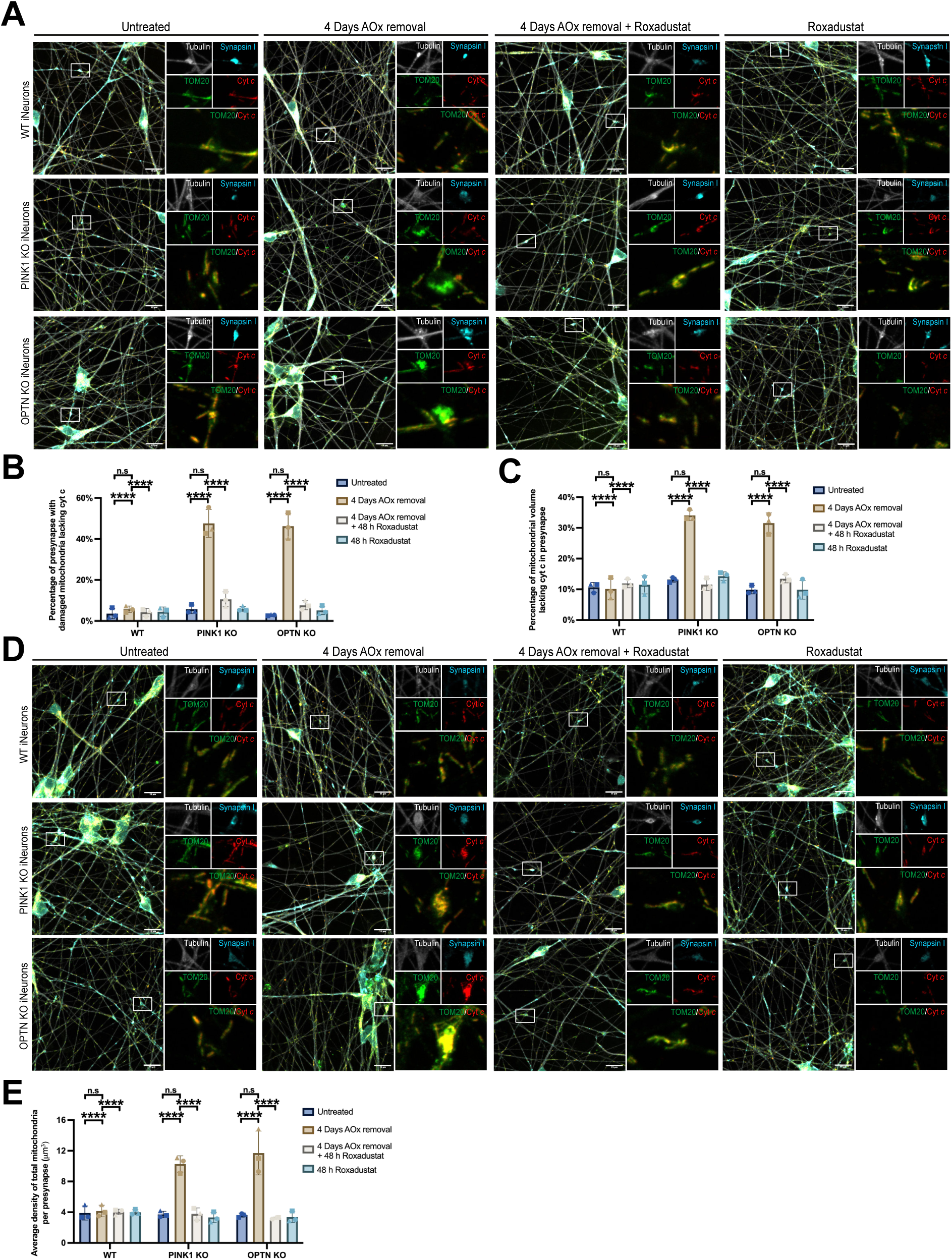
Roxadustat restores mitochondrial health and presynaptic mitochondrial density in PINK1/Parkin-mitophagy defective neurons and prevents cytochrome *c* leakage. (A-E) WT, PINK1 KO and OPTN KO iNeurons were treated pan-caspase inhibitor Q-VD-OPh (QVD) and cultured with either AOx removal media for 4 days or a combination of AOx removal for 4 days with 40 µM Roxadustat treatment in the final 48 h. The neurons were immunostained for Synapsin I, TOM20, Cytochrome *c* (Cyt *c*) and Tubulin (A and D). The percentage of mitochondrial volume lacking cyt *c* (B), the percentage of the number of presynapse with damaged mitochondria lacking cyt *c* (C) and the average density of total mitochondria per presynapse (E) were quantified. Data in (B), (C) and (E) are mean ± SD from three independent experiments. 30-60 presynapses were analysed per sample for each independent experiments. Two-way ANOVA. ****p<0.0001. n.s., not significant. Scale bars, 15 µm.

**Figure S10.**
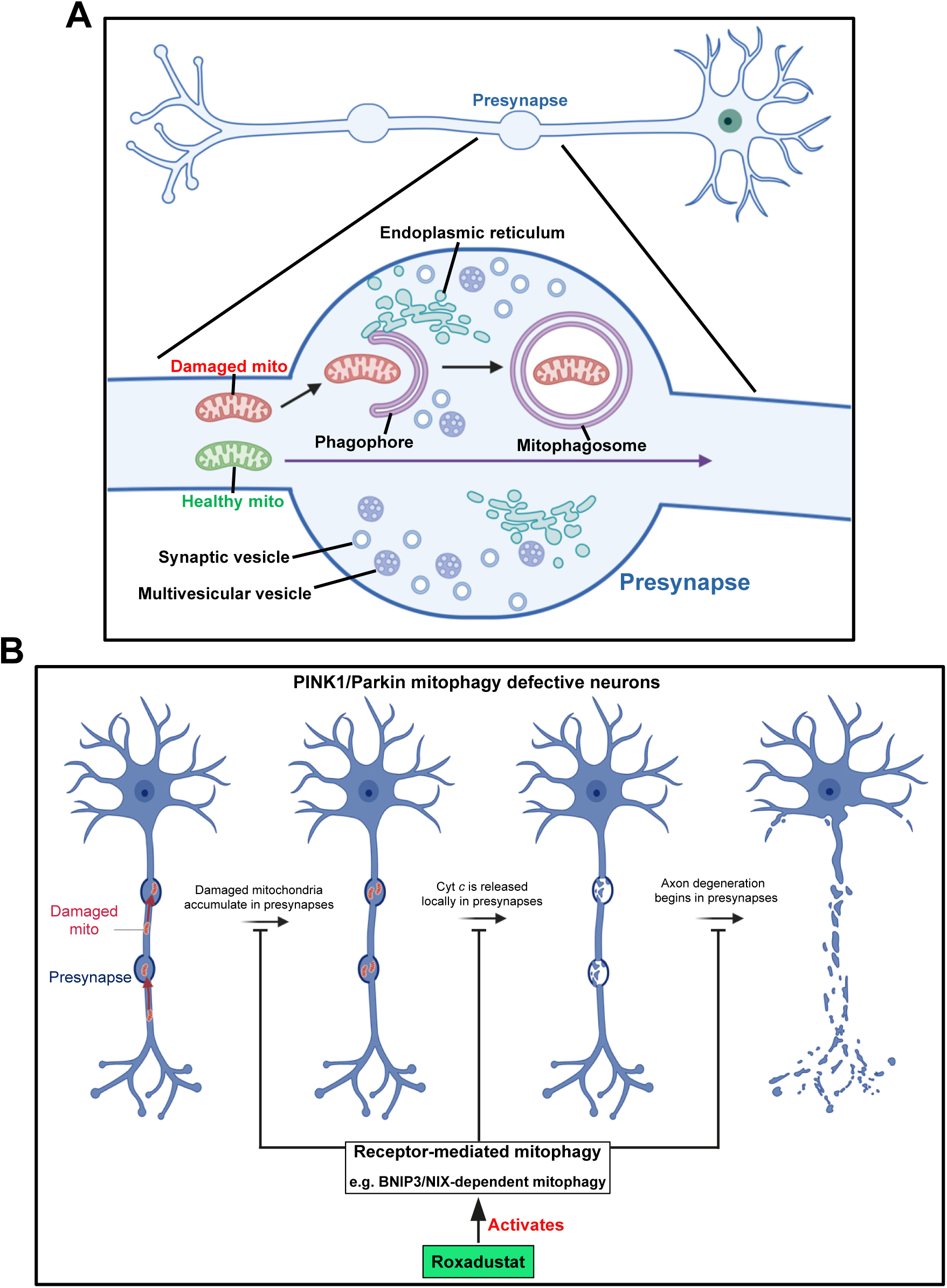
Presynapses are pit stops for PINK1/Parkin mitophagy and Roxadustat activates receptor mediated-mitophagy to prevent initiation of axon degeneration at presynapses in PINK1/Parkin defective neurons. (A) Schematic of the presynapse acting as a pit stop for PINK1/Parkin mitophagy to capture damaged mitochondria. (B) A model for the mechanism of Roxadustat preventing damaged mitochondria-induced axon degeneration in PINK1/Parkin-mitophagy defective neurons.

## Video descriptions

(Video S1. AIVE_FIB-SEM_reslice, related to Figure S4A.avi)

**Video S1.** FIB-SEM slices (top) paired with the corresponding AIVE results (bottom) in the XY projection, related to Figure S4A.

The image stack is displayed progressively through the slices. The stack is then resliced to the XZ direction and sliced through again. Scale bar = 2000 nm.

(Video S2. AIVE_CombinedStacks_Synced, related to Figure 4 and S5.avi)

**Video S2.** The AIVE result paired with the FIB-SEM slices, related to Figure 4 and S5.

The video shows a cut through of the AIVE reconstruction with the corresponding FIB-SEM stack slices. The images are shown in the XY projection.

(Video S3. AIVE_Cutthrough, related to Figure 4C and S5B.avi)

**Video S3.** Animated 3D rendering from Figure 4C and S5B, showing the presynapse with three subset areas of interest, related to Figure 4C and S5B.

The AIVE result for the presynapse area was aligned to the FIB-SEM stack in Blender. The main structure (orange) represents all membranes in proximity to the synapse. The structure is cut through from above in the XZ plane. As the video progresses the location of the three subset areas of interest are revealed in situ.

(Video S4. AIVE_MP3, related to Figure 4D.avi)

**Video S4.** Animated rotation of the 3D rendering from Figure 4D, showing an almost complete mitophagosome reconstructed with AIVE, related to Figure 4D.

This structure is aligned to an OPTN positive puncta by FIB-SEM CLEM. The reconstructed cube is 1000×1000×1000 nm. The video begins with all structures displayed. The plasma membrane (gold) is peeled back, followed by the mitochondria (red), and an adjacent multilamellar body (teal). The ER (dark blue) is then removed to reveal the shape of the almost complete mitophagosome (green). By hiding the internal mitochondria we can appreciate the complexity of the phagophore shape in 3D. The internal mitochondria are revealed to include a cluster of three mitochondrial fragments. The phagophore is then displayed again along with the ER within 100 nm of the phagophore surface.

(Video S5. AIVE_MP2, related to Figure S5C.avi)

**Video S5.** Animated rotation of the 3D rendering from S5C, showing the second mitophagosome associated with an OPTN puncta is reconstructed with AIVE, related to Figure S5C.

This area cube is also 1000×1000×1000nm and the video begins with all structures displayed. The plasma membrane (gold) and ER (dark blue) are removed to reveal several phagophore membranes (green) and mitochondria (red). The area also contains a golgi-like structure (pink). The neighbouring structures are then removed to focus on the main mitophagosome. This structure is rotated to reveal a small opening is still present in the phagophore membrane. The internal mitochondria are revealed alone. Finally, the structure is displayed along with the ER located within 100 nm of the phagophore surface.

(Video S6. AIVE_MP1, related to Figure S5D.avi)

**Video S6.** Animated rotation of the 3D rendering from S5D, showing an early phagophore forming near an intact mitochondrion reconstructed with AIVE, related to Figure S5D.

The reconstructed area measures 600×600×1000 nm. The video begins with all structures displayed. The ER (dark blue) and the other nearby mitochondria (red) are removed to reveal the main mitochondria and the phagophore (green). An additional structure is present on the middle of the mitochondrion that may represent a mitochondrial derived vesicle (magenta). The phagophore is then displayed alone before revealing the ER within 100 nm of the phagophore surface.

(Video S7. AxonSegmentation_4k, related to Figure 4F.avi)

**Video S7.** Animated 3D rendering of the presynapse of interest and its surrounding neurites, related to Figure 4F.

The AIVE result was precisely segmented into the presynapse structure (orange) and surrounding neurites (cyan). Three contacting dendrites were individually segmented (purple) with dendritic spines visible on each. The complexity of the neurites and their contact points are best appreciated in 3D.

## Materials and Methods

### Human induced pluripotent stem cells (iPSCs)

#### Culturing and Passaging iPSCs

Human WTC11 iPSCs expressing doxycycline-inducible NGN2 expression cassette in AAVS1 locus and pC13N-dCas9-BFP-KRAB targeted to human CLYBL locus were cultured in Essential 8 Medium with E8 supplement (E8 medium) (Thermo Fisher Scientific; A1517001) on Matrigel (Falcon Corning; 356231) coated vessels.^90–92^ The Matrigel coating was done by diluting the stock Matrigel 1:50 with KnockOut DMEM (Thermo Fisher Scientific; 10829018) and incubating at 37°C overnight. iPSCs were cultured at 37°C and 5% CO_2_ cell culture incubator and full medium change was performed every two days. Cells were passaged once they reached 70-80% confluency. Cells were washed twice with DPBS (Thermo Fisher Scientific; 14190144), incubated with 1:5 diluted Accutase (Thermo Fisher Scientific; A1110501) in DPBS for 5 minutes at 37°C. Once detached, cells were resuspend in E8 media and centrifuge in 15 ml falcon tube for 5 minutes at 300x g at room temperature. The cell pellet was then resuspended in E8 medium supplemented with 10 mM ROCK inhibitor (Hello Bio; Y-27632) and plated out as required onto a pre-coated Matrigel plate.

#### Differentiating iPSCs into i3Neurons

The standard established protocol was used to differentiate WTC11 iPSCs into cortical-like i3Neurons.^43,92^ Cells were maintained at 37°C 5% CO_2_ in standard tissue culture incubators. Once the iPSCs reached 70-80% confluency, cells were detached using Accutase and pelleted as described above. The pelleted cells were resuspended in pre-differentiation neuronal media, composed of KnockOut DMEM/F12 (Thermo Fisher Scientific; 12660012) supplemented with 1X N2 supplement (Thermo Fisher Scientific; 17502048), 1X MEM Non-Essential Amino Acids Solution (NEAA) (Thermo Fisher Scientific; 11140050), 1X GlutaMAX Supplement (Thermo Fisher Scientific; 35050061), 1mg/ml Laminin (Thermo Fisher Scientific; 23017015) and freshly added 10 mM ROCK inhibitor and 2 mg/ml Doxycycline (Sigma, D9891-5) to initiate differentiation. iPSCs were plated 4×10^5^ cells per Matrigel-coated well of a 6-well plate in 2 ml of pre-differentiation neuronal media. After 24 and 48 hours, media was replaced with pre-differentiation neuronal media without ROCK inhibitor. After 72 hours, the pre-differentiated iPSCs were washed twice with DPBS, lifted using Accutase and spun down as described above. The pre-differentiated iPSCs were resuspended in Bambanker (NIPPON Genetics; BB02) for - 80°C storage or resuspended in Classic Neuronal Media (CNM) with freshly added 10 mM ROCK inhibitor and 2 mg/ml Doxycycline and plated on poly-D-lysine coated vessels for neuronal maturation. CNM consisted of 50% Neurobasal (Thermo Fisher Scientific; 21103049) and 50% KnockOut DMEM/F12 supplemented with 1X N2, 1X NEAA, 0.5X GlutaMAX, 10 ng/ml Brain-Derived Neurotropic Factor (BDNF) (reconstituted in 0.1% BSA/DPBS) (PeproTech; 450-02), 10 ng/ml Neurotrophin-3 (NT-3) (reconstituted in 0.1% BSA/DPBS) (PeproTech; 450-03), 0.5X B-27 supplement (Thermo Fisher Scientific; 17504001), and 1mg/ml Laminin. Pre-differentiated cells were seeded in 35 mm m-Dish with grids (ibidi; 81166), 6-well plate or 8-well chamber slide for FIB-SEM CLEM experiment, Western Blotting and fixed or live cell imaging respectively. The seeding density was 3×10^4^ cells/cm^2^ of growth area. Biweekly half media changes were performed to maintain the cultures. All experiments were performed 10 days after the pre-differentiation stage (DIV 10), unless otherwise specified. All the iPSCs used for this study were differentiated before passage 24.

### Human embryonic stem cells (ES cells)

#### Culturing and Passaging ES cells

The WT, PINK1^-/-^ and OPTN^-/-^ human ES cells (H9, WiCell Institute) used in this study were obtained from Harper’s lab.^93,94^ All these ES cell lines were expressing doxycycline-inducible NGN2 expression cassette in AAVS1 locus. These cells were cultured in E8 medium on Matrigel coated plates. The culturing and passaging procedures were identical to the iPSCs as described above, except the full medium change was performed daily instead of every two days.

#### Differentiating ES cells into iNeurons

When cells reached 70-80% confluency, they were lifted with Accutase, spun down and resuspended in ES pre-differentiation media (KnockOut DMEM/F12 supplemented with 1X N2 supplement, 1X NEAA, 10 ng/ml BDNF (reconstituted in 0.1% BSA/DPBS), 10 ng/ml NT-3 (reconstituted in 0.1% BSA/DPBS), 1 mg/ml Laminin, 10 mM ROCK inhibitor and 2 mg/ml Doxycycline. Cells were seeded at 2×10^4^/cm^2^ on Matrigel-coated plates. After 48 hours, the media was replaced with ES neuronal media composed of Neurobasal supplemented with 1X GlutaMAX, 0.5X B-27 supplement, 10 ng/ml BDNF, 10 ng/ml NT-3 and 1 mg/ml Doxycycline. On day 4 and 6, a half medium change was performed. On day 7, cells were washed twice with DPBS, detached with Accutase and spun down as above. The cell pellet was either resuspended in Bambanker for -80°C storage or resuspended in ES neuronal media with 1 mg/ml Doxycycline and 1 mM ROCK inhibitor and plated at 3×10^4^ cells/cm^2^ of growth area on Matrigel-coated vessels for the final neuronal maturation. Doxycycline and ROCK inhibitor were removed from ES neuronal meida on Day 10. iNeurons were used for experimentation on Day 14 (DIV14).

#### Generation and maturation of neurons from human iPSCs into midbrain neurons

iPSCs were maintained in mTESR1 medium (STEMCELL Technologies; 85850) on tissue culture ware coated with hESC-qualified Matrigel (Corning; 354277), as per manufacturer’s instructions. Differentiation of iPSCs into midbrain neural progenitor cells (NPCs) was performed using an embryoid body (EB)-based protocol using dual SMAD inhibition, as described previously.^57,95^ Midbrain NPCs were expanded to four passages on hESC-Matrigel coated plates in neural progenitor medium (NPM) and cryobanked in liquid nitrogen. NPM was composed of DMEM/F12 + GlutaMax™ basal medium (Thermo Fisher Scientific; 10565018) supplemented with 1× N2 (Thermo Fisher Scientific; 17502048), 1× B27 (Thermo Fisher Scientific; 17504044), FGF-8b (100 ng/mL, PeproTech; 100-25), Sonic Hedgehog (200 ng/mL, R&D Systems; 1314SH) and laminin (1 μg/mL, Thermo Fisher Scientific; 23017015). For neuronal maturation, NPCs were thawed and expanded for one extra passage in NPM. When confluent, NPCs were dissociated using Accutase (STEMCELL Technologies; 07920) and re-plated at 2×10^5^ cells/cm^2^ onto 5 µg/mL laminin-coated tissue culture plates in midbrain neural maturation medium (NMM). Midbrain NMM was composed of BrainPhys Neuronal Medium supplemented with N2A, SM1, 200 nM ascorbic acid, 1 µg/mL (1.2 nM) laminin, 20 ng BDNF (STEMCELL Technologies), 20 ng GDNF (STEMCELL Technologies) and 0.5 mM dibutyryl cyclic-AMP (Sigma-Aldrich). At this time (day 0 of maturation), seven NPC lines from three healthy individuals (two clones per individual plus one human ESC line) were mixed together in NMM to form a neuronal village model. Neuronal village cultures were maintained for 14 days following differentiation. Half media changes were performed every 2-3 days before replating onto DAP-coated coverslips ^59^ with an additional coating of 10 µg/mL poly-L-ornithine (Sigma-Aldrich) and 5 µg/mL laminin, in NMM containing half concentrations of growth factors. Cells were maintained at 37°C 5% CO_2_ in standard tissue culture incubators, and media was refreshed every 2-3 days via half medium changes, performed by Assist Plus (Integra). Cultures were matured for at least 3 months *in vitro* (90DIV) before experiments.

### Cloning and generation of stable iPSC and ES cell lines

A template PLX311 vector backbone with EF1a promoter was generated by digesting PLX311-Cas9 (Addgene: 118018) with restriction enzymes NheI and SpeI, followed by Quick CIP (New England Biolabs; NEB#M0525) treatment to prevent re-ligation of linearised plasmid DNA. Linearised PLX311 vector without the Cas9 gene was then purified via DNA gel purification. Open reading frames (ORFs) of mito-EBFP2 (a gift from Michael Davidson; Addgene #55248), EGFP-OPTN (Addgene #188784), mt-Keima (a gift from Richard Youle; Addgene #72342), mito-mKeima^XL^ (a gift from Wade Harper^96^) were first PCR amplified. An extra 13 bp sequence (catcgattgatca) was inserted before the ORFs to make the expression more stable. The PCR products were introduced into linearised PLX311 template vector using NEBuilder® HiFi DNA Assembly Cloning kit (New England Biolabs). All constructs were sequence-verified.

To generate stably transfected iPSC or ES cell lines, lentiviral vectors (PLX311-mito-EBFP2, PLX311-EGFP-OPTN, PLX311-mt-Keima, PLX311-mt-Keima-xL) were transfected in HEK293T together with lentiviral packaging plasmids (VSV-G and psPAX2) using Lipofectamine LTX (Life Technologies) for 16 hours. Transfection media was replaced with fresh pre-warmed Dulbecco’s Modified Eagle Medium (DMEM) (Thermo Fisher Scientific; 11885084) supplemented with 10% (v/v) Fetal Bovine Serum (Moregate; Batch 93301104), 4.5 g/L glucose, 1X GlutaMax, 1X NEAA, 10 mM HEPES and 50 units/ml Penicillin and 50 μg/ml Streptomycin (Thermo Fisher Scientific; 15140122) to allow assembly and secretion of lentiviruses. After 24 hours incubation, supernatant containing lentivirus was collected, filtered, and mixed with Lentivirus Precipitation Solution (ALSTEM; VC100). The lentivirus mixture was vortexed, then incubated at 4°C on a rocking platform mixer at 30 rpm for 24-48 hours. Therafter, the lentivirus mixture was centrifuged for 1 hour at 1600x g at 4°C before the lentivirus pellet was resuspended in E8 medium with 10 mM ROCK inhibitor and 1 mg/ml polybrene. The resuspended lentivirus medium was then added to iPSC or ES cells that had been seeded 1 hout prior onto Matrigel coated plates. After 24 hours, the lentivirus medium was replaced with fresh E8 medium containing 10 mM ROCK inhibitor. Transduced cells were expanded in E8 medium without ROCK inhibitor for at least two passages before differentiation.

### Mitophagy induction treatment

#### AOx removal-induced mitophagy for i3Neurons and iNeurons

Antioxidant (AOx) removal approach was used for inducing neuronal mitophagy via promoting mild oxidative stress.^42^ A half media change in the neuronal media (CNM or ES neuronal media) was performed two days prior to antioxidant removal. For the treatment, neuronal media was fully replaced with fresh control or AOx-free neuronal media in which the B-27 supplement was substituted with B27 supplement minus antioxidants (Thermo Fisher Scientific; 10889038). This was followed by a two-thirds media change immediately to ensure complete washout of the original media. Different time points are indicated in figure legends. For long term AOx removal (4 days or more), a half media change was performed every two days.

#### AOx removal-induced mitophagy for iPSCs-derived midbrain neurons

For mitophagy induction, 157DIV midbrain iPSC-derived neuronal cultures underwent AOx removal. A half media change in midbrain NMM was performed two days prior to AOx removal. On the day of the 6 hours AOx treatment, a full medium change was performed using AOx-free medium comprising BrainPhys basal with AOx-free B27, N2A, BDNF, GDNF, cAMP and laminin. Two-thirds of the media was immediately replaced with fresh AOx-free medium to ensure complete washout of the original medium. Untreated cells received the same medium changes except with AOx-containing medium (BrainPhys basal with B27, N2A, BDNF, GDNF, cAMP, laminin and ascorbic acid).

#### Roxadustat-induced mitophagy for iNeurons

Roxadustat (R&D System; 808118-403) was used for inducing BNIP3/NIX-dependent mitophagy via activating HIF1α signalling through inhibiting prolyl-4-hydroxylase activity. A half media change in the ES neuronal media was performed two days prior to the treatment. On the day of the treatment, half of the media was removed and replaced with fresh ES neuronal media containing 80 µM Roxadustat, making the final Roxadustat concentration 40 µM. In the condition where the neurons were pretreated with AOx-free neuronal media for two days, the same procedure was done except half of the media was replaced with fresh AOx-free neuronal media containing 80 µM Roxadustat.

### Live cell imaging

For live cell imaging, both iPSCs-derived i3Neurons and ESCs-derived iNeurons were seeded in ibiTreat, polymer bottom 8-well chamber µ-slide (Matrigel coated) and cultured in phenol-red free CNM and ES neuronal media, respectively. All live cell images were acquired on the Leica Stellaris 8 confocal microscopy with FLIM (using lighting mode) under 40x/1.3 numerical aperture (oil immersion, HCPL, APO, CS2, Leica Microsystems) objective lens with 2.5X zoom. The microscope was equipped multiple Leica Power HyD detectors and a thermally regulated incubator hood with CO_2_ controller (37°C and 5% CO_2_). Uniform laser intensity and exposure duration were consistently applied across all samples, with adjustments to brightness and contrast standardized using identical minimum and maximum display values in ImageJ/Fiji.

#### Detecting mitochondrial membrane potential

WT i3Neurons expressing mito-EBFP2 were cultured. Samples were stained prior to imaging for 20 minutes at 37°C with 500 nM tetramethylrhodamine methyl ester (TMRM) and 500 nM SiR-Tubulin (Cytoskeleton: CY-SC002). All images were acquired in three-dimensional (3D) by optical sectioning with a minimum z stack range of 3.6 μm and a maximum voxel size of 56.7 nm laterally (*x*, *y*) and 517 nm axially (*z*) with a scan speed of 700 Hz. The max-projected deconvoluted images were displayed. The depolarised mitochondria are indicated by the lack of TMRM staining (i.e. mito-EBFP2 positive but TMRM staining negative).

#### Mt-mKeima^XL^ mitophagosome-lysosome fusion mitophagy measurement

iNeurons expressing mt-mKeima^XL^ were stained with 1 µg/ml Hoechst33342 and 500 nM SiR-Tubulin for 20 minutes at 37°C prior to imaging. The Hoeschst33342 and SiR-Tubulin were excited using 405 nm and 652 nm laser lines. For ratiometric imaging of the mitophagy reporter mt-mKeima^XL^, a sequential 488 nm and 552 nm laser excitation with scan speed of 700 Hz was used, and the fluorescence emission was collected between 577-708 nm. The acidified lysosomal mt-mKeima^XL^ was identified by the longer-wavelength (552 nm) excitation being predominated. All images were acquired in 3D by optical sectioning with a minimum z stack range of 1.05 μm and a maximum voxel size of 42 nm laterally (*x*, *y*) and 353 nm axially (*z*). The max-projected deconvoluted images were displayed. Six images were captured per condition and three independent experiments were performed. All the images were randomised and blinded prior to analysis. The number of mitolysosomes (acidified mt-Keima^XL^) was counted semi-manually with the assistance of the “Find Maxima” function in ImageJ/Fiji.

#### Mitochondrial ROS measurement

WT iNeurons were stained with 100 nM Mitotracker Green FM (MTG), 0.5 µM MitoSOX red and 500 nM SiR-Tubulin for 20 minutes at 37°C prior to imaging. The concentration of MitoSOX red used in this study was optimised to ensure untreated neurons exhibited minimal MitoSOX red intensity. All images were acquired in 3D by optical sectioning with a minimum z stack range of 1.05 μm and a maximum voxel size of 57 nm laterally (*x*, *y*) and 353 nm axially (*z*) with the scan speed of 400 Hz. The max-projected images were displayed. Six images were captured per condition and three independent experiments were performed. To measure the ratio of MitoSOX red to MTG, first, the area of the mitochondria in MTG and MitoSOX red channels were calculated. Image thresholding was applied using gate settings of (25, 150) for the MTG channel and (5, 100) for the MitoSOX red channel. These binary images were subsequently analysed using the “Analyse Particles” function in ImageJ/Fiji to quantify the area occupied by mitochondria in each channel. Finally, the ratio of MitoSOX red to MTG channel areas occupied by mitochondria was determined. The higher the ratio of MitoSOX red to MTG, the higher the amount of mitochondrial ROS the mitochondria produced.

### Immunoblotting

To prepare samples for Western blot analysis, cells from the 6-well plate were harvested in ice-cold 1X DPBS using cell scrapers and lysed in 1X lithium dodecyl sulfate (LDS) sample buffer (Life Technologies, NP0007) supplemented with 100 mM dithiothreithol (DTT) (Astral Biosciences, 3483-12-3). For the Caspase 3/Cleaved caspase 3 experiment shown in Figure 7C, the neuronal media were collected before harvesting in the ice-cold 1X DPBS (supplemented with 500 µM phenylmethylsulfonyl fluoride (PMSF) protease inhibitor). After the neuronal media were spun down, the dead cell pellets were combined with the live cell pellets collected from scraping, and lysed together in 1X LDS sample buffer with DTT. For the PINK1 immunoblotting experiment shown in Figure S2E, cells were lysed in 1X SDS sample buffer [5% (w/v) SDS, 10% (v/v) glycerol, 100 mM DTT, and 50 mM tris-Cl (pH 6.8)]. All the samples were heated at 99°C with shaking for 10 minutes. Approximately 20-80 µg of protein per sample was analysed by NuPAGE Novex 4 to 12% bis-tris gels (Life Technologies) according to the manufacturer’s instructions and electro-transferred to polyvinyl difluoride membranes (PVDF) and immunoblotted using indicated antibodies (see Key Resource Table for the antibodies used in this study).

### Focus ion beam scanning electron microscopy (FIB-SEM) with live-correlative light-electron microscopy (live-CLEM) and Artificial Intelligence directed Voxel Extraction (AIVE)

#### Sample preparation and imaging

35 mm ibidi µ-dishes with a polymer coverslip bottom and an imprinted 500 µm cell location grid were used for FIB-SEM with live-CLEM experiment. The ibidi µ-dishes were glow discharged with the PELCO easiGlow Glow Discharge Cleaning System before coating with Poly-D-Lysine. iPSC-derived i3Neurons expressing EGFP-OPTN were seeded onto the coated 35 mm ibidi dish and the neurons were matured in phenol-red free CNM. On the day of the experiment, the i3Neurons were stained with 1 µg/ml Hoechst 33342, 500 nM TMRM and 500 nM SiR-Tubulin for 20 minutes at 37°C prior to imaging. The Leica SP8 confocal microscopy with 63x/1.4 numerical aperture (oil immersion, HCPL, APO, CS2, Leica Microsystems) objective lens was used for this experiment. The microscope was equipped with a Leica Power HyD detector, two PMT detectors and a thermally regulated incubator hood with CO_2_ controller. After locating the neuron or area of interest, the grid position was identified. An overview image in 3D was acquired using optical sectioning with a minimum z stack range of 6 µm, and a maximum voxel size set to 180 nm laterally (x, y) and 838 nm axially (z). This was an essential step for relocating the neurons for CLEM. A zoomed-in 3D image with a smaller Z-step of 334.5 nm axially was subsequently acquired. Concurrently pre-warmed 4% (v/v) paraformaldehyde (PFA) in 0.1 M phosphate buffer was added to the ibidi µ-dish. Once the optical image acquisition was completed, PFA fixation was left to continue at room temperature for 30 minutes. The sample was then post-fixed with 2.5% glutaraldehyde in 0.1 M sodium cacodylate (NaCac) buffer (pH 7.4) for 24 hours at 4°C, then rinsed twice with 0.1 M NaCac buffer for tertiary fixation and contrasting. All subsequent stages were microwave assisted using a BioWave Pro microwave system (Pelco). Tertiary fixation began with 2% osmium tetroxide (OsO_4_) and 1.5% Potassium ferricyanide (K3Fe(III)(CN)6) in 0.1 M NaCac buffer on ice for 1 hour, then crosslink samples with 1% (w/v) Thiocarbohydrazide in water, and followed by the final fixation with 1% OsO_4_ in water. Samples were washed with MilliQ water in between each step, and three microwave duty-cycles (120 s on, 120 s off) at 100 W under vacuum was used at each stage. The samples were then *en bloc* stained with 2% (w/v) aqueous uranyl acetate^97^, followed by Walton’s Lead Aspartate staining ^98^. Three microwave duty-cycles (120 s on, 120 s off) at 100 W under vacuum for both staining processes. Relocate the neuron targets using Leica M125 dissection microscope with an IC80 HD camera (Leica Biosystems) and mark the target area. Perform a microwave assisted dehydration by graduated ethanol series (80%, 90%, 95%, 100%, 100% (w/v); each at 150 W for 40 s) and propylene oxide (100%, 100% (w/v); each at 150 W for 40 s). The samples were infiltrated with Araldite 502/Embed 812 by graduated concentration series in propylene oxide (25%, 50% 75% 100%, 100% (v/v); 180 s at 250 W under vacuum), then polymerized at 60°C for 24 hours. The target area from the resin-embed samples were relocated and marked using dissection microscope (Leica Biosystems), then cut out from the ibidi µ-dish and mounted onto 3.18 mm diameter aluminium rods with Araldite 502/Embed 812, and polymerised for an additional 48 h at 60°C. Excess polymer-coverslip and resin was removed from the sample to create a square block face using an Ultracut UCT ultramicrotome (Leica Biosystems) with a freshly made glass knife. The sides of the resin block were coated with silver pasta and cure at 93°C for 2 hours. 170 µm of polymer-coverslip from the square block face was removed using the Ultra UCT ultramicrotome (Leica Biosystems) equipped with a glass knife, leaving around 20 µm of polymer coverslip. Samples were gold coated using Leica ACE200 for 3 minutes and the precise thickness of the polymer was measured using the cryo-Helios G4 UX FIB-SEM (FEI). Samples then removed from the FIB-SEM and final ultrafine trimming was performed using Ultra UCT ultramicrotome (Leica Biosystems) equipped with a diamond knife (Ultra 45° Diatome). The final polymer thickness on the block face needs to be within 3 – 4 µm thick. The block face of the samples was cleaned with high pressure nitrogen gas, and two images were acquired using Leica M125 dissection microscope with an IC80 HD camera (Leica Biosystems) at 7.5X and 10X magnification and imported into the MAPS (v2.5; FEI) program to enable sample navigation during FIB-SEM imaging. Angled gold coating was performed twice using Leica ACE200 for 5 minutes. The samples were imaged using a cryo-Helios G4 UX FIB-SEM (FEI) operating at 3 nm per pixel (2 kV, 100 pA, HFV 18.44 μm, 4 μs dwell time), with ion milling at 10 nm per slice (Gallium, 30 kV, 2.4 nA) using the Auto Slice And View (v4.1; FEI) software.

#### FIB-SEM image alignment

The FIB-SEM stack was initially cropped to remove excess empty area and gold coating before spatial registration using the ImageJ (v1.53t) Register Virtual Stack plugin, with affine feature extraction and rigid registration set to the last frame. Settings were run default with no interpolation. The resulting transformed image was cropped to an area of 3072×1024×601 voxels around the bouton. This processed stack formed the working image for the CLEM alignment as well as the three arms of the AIVE image analysis pipeline (below).

#### FIB-SEM CLEM

Orthoslices (XZ) of the aligned FIB-SEM data were spatially aligned to the deconvolved optical data using established procedures for CLEM alignment. The stack was initially rotated 5 degrees in Y in the XY projected angle to align the dish to 0 degrees. The stack was then resliced in YZ with no interpolation. The stack was then rotated again by 7 degrees in Z to align the dish to 0 degrees. The image was then resliced to XZ to match the confocal imaging plane. A final rotation of 8 degrees in X was performed to complete the alignment. This adjusted stack was used for CLEM orientation and location of OPTN positive structures only.

#### AIVE reconstruction

The aligned working image FIB-SEM stack was reconstructed in 3D using an adapted version of AIVE (Artificial Intelligence directed Voxel Extraction)^52^. This involved the following Parallel image analysis pipeline:

##### A: Organelle segmentation maps

The working image was examined for areas of interest. These areas were selected based on overlap with OPTN signal by FIBSEM-CLEM. These areas were subset and segmented based on organelle type. Semi-automatic thresholding and manual curation of structures of interest was performed using Microscopy Image Browser software. Each segmented structure was exported as a binary .tif map. Resulting binary maps were processed to account for uncertainty between binary slices using ImageJ macro: AIVE_1.ijm. This included an initial 10nm gaussian blur. As this reduces the signal intensity, the original binary was then added back onto the result using ImageJ image calculator, followed by a second 10nm gaussian blur. This image was used to segment the organelles on a per structure basis for the final AIVE output.

##### B: Image source

The working image was pre-processed with a low pass noise filter (Average result between the original working image and a 3nm gaussian blur) followed by contrast limited adaptive histogram equalisation (CLAHE). To approximate the CLAHE filter in 3D the working image was filtered in each of the 3D dimensions (XY, XZ, YZ) using the 2D ImageJ CLAHE filter and the average result designated as the final “Source” image.

##### C: AI membrane prediction

Microscopy Image Browser v2.81 was used to generate sparse labels for the categories “Resin”, “Cytoplasm”, “Matter” and “Membrane” from a small image subset. Dense labels were generated by curating the output of this initial model, trained with WEKA software v3.8.5, and applied across the full training subset (slice 401 of the working image stack). 50,000 voxels per class, representing 1.59% of total voxels of this slice, (200,000 total voxel instances) were then used to train the final AI prediction model. A custom macro harnessing the Fiji Trainable WEKA Segmentation plugin was used to generate 170 2D and 3D image “features” describing the local environment of each voxel. Results were encoded as 32-bit images and an adapted macro was used to convert these instances to an .arff file for import to the WEKA software. A random forest machine learning model was trained from these data using 10 folds of cross validation, a bag size of 10%, batchsize=100, No random break ties, OutOfBag=True, AttributeImportance=True, with an unlimited tree depth, 9 random features, 200 iterations per round and a starting seed of 1337. The final model was accessed from a second custom macro to output class predictions for the full working image as a 32-bit image with values between 0 and 1. Model self-accuracy to the dense labels was 97.482%. Only the result for the membrane class was used for the AIVE merge.

#### D: AIVE merge

The Image B: Image Source was converted to 8-bit and multiplied by each of the organelle segmentation maps from Image A. Image C: The membrane prediction, was also converted to 8-bit then multiplied by each of the blurred segmentation maps followed by resetting the range back to 8-bit and filtering with a 10nm gaussian blur. The output of the source and membrane prediction results were then averaged to create a consensus that was taken as a threshold above 56. The .tiff result was imported to Paraview v5.9 and contoured using the consensus value. Contours were then exported as .3dx files and imported to Blender v3.0.0 for animation. Software analysis of images was performed using an Intel® Core™ i9-10980XE workstation with 64GB of RAM. Animations were rendered using the WEHI High Performance Computing cluster using a batch script.

### Immunocytochemistry

#### Immunofluorescence assay (IFA) for iPSC-derived i^3^Neurons and ESC-derived iNeurons

For immunofluorescence, both iPSCs-derived i^3^Neurons and ESCs-derived iNeurons were cultured in ibiTreat, polymer bottom 8-well chamber µ-slide (Poly-D-Lysine or Matrigel coated). Upon fixation, half of the neuronal media were replaced with pre-warmed (37°C) fixative and fixed for 15 minutes at room temperature. The fixative was freshly made with 4% PFA in 2X microtubule stabilisation buffer (160 mM PIPES (pH 6.8), 10 mM EGTA, 2 mM MgCl_2_, H_2_O) and DPBS. Samples were gently washed three times with DPBS, permeabilised with 0.1% (v/v) Triton X-100 in DPBS for 10 minutes, and then blocked with blocking buffer (3% (v/v) goat serum in 0.1% Triton X-100/DPBS) for 20 minutes. Samples were incubated with indicated primary antibodies diluted in blocking buffer for 1.5 hours at room temperature. The only exception was the p-UB (ser65) staining, which was incubated overnight at 4°C on a rocker. Samples were washed three times with DPBS then incubated with secondary antibodies conjugated to AlexaFluor-405, AlexaFluor-488 or AlexaFluor-555 that were diluted in blocking buffer for 1.5 hours. Samples were washed three times with DPBS then incubated with AlexaFluor 647 conjugated β-Tubulin (Cell Signaling Technology, 3624S) diluted in blocking buffer for 2 hours. Final three times DPBS washes were performed prior to adding 150 µl of mounting media into the well. All primary and secondary antibodies used in this study are listed in the Key Resources Table.

Imaging of the samples was done with the Leica Stellaris 8 confocal microscopy with FLIM under 40x/1.3 numerical aperture (oil immersion, HCPL, APO, CS2, Leica Microsystems) objective lens with 2.5X zoom. For the cytochrome *c* release and the mtDNA release experiments, 6.5X zoom images were acquired and shown as the inset images. Images were acquired in 3D by optical sectioning with a minimum z stack range of 2.1 μm and a maximum voxel size of 56.7 nm laterally (*x*, *y*) and 353 nm axially (*z*) with the scan speed of 400 Hz or 700 Hz (for the lighting mode). All images displayed as max-projection. Except for FigureS7A, max-projection of deconvoluted images were shown. For image analysis, 6-8 images were taken per sample at random stage positions, and 30-60 presynapses were analysed per sample.

#### Mitophagy experiment of iPSC-derived midbrain neurons

Cells were fixed using pre-warmed (37°C) 4% paraformaldehyde in 2X microtubule stabilisation buffer for 15 minutes, followed by three DPBS washes. Fixed cells were permeabilised in permeabilisation buffer (0.1% Triton X-100 in DPBS) for 10 minutes then replaced with blocking buffer (DPBS containing 0.1% Triton X-100 and 3% goat serum) for 20 minutes. Thereafter, cells were incubated with primary antibodies diluted 1:500 in blocking buffer for 1.5 hours rocking at room temperature. Primaries were anti-mouse WIPI2 (Abcam; ab105459), anti-chicken HSP60 (Novus; NBP3-05536), and anti-rabbit Synapsin I (Novus; NB300-104). Cells were washed three times with DPBS then incubated with secondary antibodies diluted in blocking buffer, for 1.5 hours at room temperature. Secondaries were AlexaFluor (AF) 555 goat anti-mouse (Thermo Fisher Scientific; A21422) 1:500, AF488 goat anti-chicken (Thermo Fisher Scientific; A11039) 1:500, and AF405 goat anti-rabbit (Thermo Fisher Scientific; A31556) 1:250. Cells were washed three times with DPBS then incubated with conjugated antibody staining buffer comprising β-Tubulin (9F3) Rabbit mAb (AF647 Conjugate) (CST; 3624S). After 2 hours incubation at room temperature cells were washed a final three times with DPBS. Coverslips were then mounted onto slides using mounting media and sealed once dried.

#### Characterisation of iPSC-derived midbrain neurons

After 90DIV or 120DIV, cells were fixed in 4% paraformaldehyde for 10 mins followed by three washes with PBS and one wash with Tris-buffered saline (TBS). Cells were permeabilised using 0.1% Triton X-100 treatment and blocked using 3% Donkey serum in 0.1 mM TBS for 60 minutes at room temperature. Primary antibody solutions were prepared in TBS++ (TBS, 0.1% Triton X-100 and 3% donkey serum) against MAP2 1:2000 (Abcam; ab5392), LMX1A (Abcam; ab139726), FOXA2 1:50 (In Vitro Technologies; RDSAF2400SP), GIRK2 1:200 (Abcam; ab65096), Calbindin 1:150 (Abcam; ab108404) and TH (R&D Systems; MAB7566). Cultures were incubated with the primary antibody solutions overnight at 4°C. Primary antibody solutions were aspirated, and cells were washed thrice with TBS at room temperature. To block, TBS++ was added to cell cultures and incubated for 1 hour at room temperature. AF-conjugated secondary antibody diluted 1:250 in TBS++ were then added and incubated for 1 hour at room temperature. Secondaries were donkey anti-mouse AF647 (Abcam; AB150111), donkey anti-chicken Alexa-Fluor 488 (Stratech Scientific APAC; 703-545-155) and Donkey Anti-Rabbit IgG AF568 (Abcam; AB175692). Cells were stained with 4ʹ,6-diamidino-2-phenylindole (DAPI nuclear stain: 1:1000; Sigm; D9542,) for 5 minutes following secondary antibody aspiration and washed thrice with PBS immediately prior to imaging. Images were acquired using an Operetta CLS™ High Content Imaging System (PerkinElmer). Twenty-five fields of view were captured per well (20X magnification) each comprising ten Z stacks every 1 µm. Custom Harmony 4.9 (PerkinElmer) pipelines were used for analysis of maximum projection images whereby each well contained at least 3000 DAPI+ nuclei.

### Whole-cell patch clamping

Coverslips containing midbrain neurons were transfected with a Synapsin:GFP lentiviral vector before being transferred into a recording chamber that was perfused with artificial cerebrospinal fluid (ACSF) at room temperature (21-23°C) using the Minipuls 3 (Gilson). Prior to perfusion, the osmolality of ACSF was tested using a Fiske Micro-Osmometer (model 210, Advanced Instruments) to ensure it was within the correct range (300-310mOsm). ACSF contained: 121 mM NaCl, 4.2 mM KCL, 1 mM MgSO_4_, 29 mM NaHCO_3_, 0.45mM NaH_2_PO_4_, 0.5 mM Na_2_HPO_4_, 1.1 mM CaCl_2_ and 20mM glucose (all chemicals from Sigma-Aldrich) and was continuously bubbled with 95% O_2_ and 5% CO_2_. Whole-cell patch clamp recordings were performed underneath an upright Olympus BX51 microscope, a 40x water-immersion objective lens and a PCO.Panda 4.2 camera. A cool-LED pE300 illumination unit at 460nm was used to visualise neurons expressing Synapsin:GFP. Borosilicate glass filaments (BF100-50-7.5, Sutter Instrument) were pulled with an open resistance of 3-6 MΩ using a P-1000 micropipette puller (Sutter Instruments). Patch pipettes were filled with an internal solution that has a pH and osmolality similar to regular physiological conditions (pH 7.3, 290-300mOsm). Intracellular solution contained: 130 mM K-gluconate, 6 mM KCl, 4mM NaCl, 10 mM Na-HEPES, 0.2 mM K-EGTA, 0.3 mM Na-GTP, 2 mM Mg-ATP, 0.2 mM cAMP, 10 mM D-glucose, 0.15% biocytin and 0.06% rhodamine.

Whole-cell patch clamp recordings were amplified using a Digidata 1550B/Multiclamp series 700B, digitised and sampled using PClamp software (v10.7 Molecular Devices) at 50k Hz for voltage-clamp recordings and 100kHz for current-clamp recordings. To record voltage-gated sodium (Na^+^) and potassium (K^+^) currents, cells were held at -70 mV in voltage-clamp mode and injected with +5 mV current steps across 20 sweeps starting from an initial drop to -75 mV. The *Maximum Nav Peak (pA)* corresponds to the sweep that had the largest sodium depolarisation in the recording. Cells were then held in C=0 to record the resting membrane potential. Action potential firing and kinetics were obtained by using a sufficient amount of current to hold the neuron at -70 mV in current-clamp mode. If holding current exceeded -100 pA that patch was considered poor quality and aborted. Neurons were subjected to a 500ms depolarising current step for 15-20 sweeps, and the *Evoked AP Frequency (Hz) (APs > -10 mV)* corresponds to the sweep that had the highest firing rate in the recording. AMPA-mediated events were recorded at -70 mV in voltage-clamp mode, while GABA-mediated events were recorded at -10 mV in voltage-clamp mode (at the reversal potentials for sodium and chloride, respectively) and they were recorded for 3 minutes with a gap-free run. All patch-clamping data was analysed using custom-built scripts and reported membrane potentials were corrected for the junction potential (∼10 mV) post analysis.

Whole-cell patch clamping exclusion criteria: To exclude unhealthy neurons and suboptimal recording conditions, only neurons with: 1) > 1 GΩ seal and 2) access resistance < 20 MΩ were included in the final analysis stages. If patch quality or cell health deteriorated such that quality control checks were failed, the patch was aborted and only recordings acquired prior to this were included.

### Confocal image analysis

#### Quantifying of the colocalisation between mitochondria and mitophagy markers

In this study we employed robust statistical methods to detect Foci on mitochondria within neuronal structures including axons and axonal boutons/presynapses. The soma and boutons/presynapses were manually segmented. Somas were excluded when analysing axons. In order to detect the Foci we implemented a combined approach utilising Robust Gaussian Fitting Library (RGFLib) Python package^99^ and HDBSCAN package^100^. RGFLib effectively differentiates peak signals from background noise through a combined approach of robust statistics for peak separation and density-based region growing for background merging. This joint segmentation/merging technique enables the precise detection of peaks of varying shapes and sizes.

To further refine the analysis, two filtering steps on the detected peaks were implemented. First, clusters exceeding or falling below a predetermined pixel size threshold were excluded, ensuring only peaks within an appropriate size range were considered for further analysis. Second, any pixel gaps within the detected peaks was filled, to ensure consistent labelling across all detected regions.

To quantify the co-localization of signals between mitochondria and p-UB (Ser65), ATG13 or EGFP-OPTN channels, a mask-based approach was employed. In this approach, the analysis result of mitochondrial channel was used as a mask, highlighting regions of interest. The analysis was carried out using RGFLib background separation method. This channel’s information was then used to mask the other channel which effectively zeroes out the intensity values in the p-UB Ser65), ATG13 or EGFP-OPTN channel outside the masked regions allowing separation of Foci peaks occurring inside mitochondria. The percentage of mitochondria that are positive for p-UB (Ser65), ATG13 or EGFP-OPTN was determined by calculating the ratio between mitochondria volume positive for foci of interest and total mitochondria volume.

#### Quantification of autophagy factor ATG9A and caspase activation marker Cleaved Caspase-3 signal intensity

To measure the intensity of ATG9A or Cleaved Caspase-3 (CC3) in specific neuronal compartment such as axons, presynapses or axon minus bouton/presynaspes, raw max-projected images were analysed using ImageJ. The somas and presynapses were manually segmented as these neuronal compartments are morphologically distinct from the axons. To assess the signal intensity in axons, somas were first excluded from the images. Image thresholding was applied to the tubulin channel. The volume occupied by tubulin was determined using the “Analyse Particles” function in ImageJ/Fiji, and the ROI was created automatically. This ROI was then applied to the ATG9A or CC3 channel, and subsequently the ATG9A or CC3 intensity per axon volume was calculated.

An extra step of analysis was needed for the ATG9A experiment. Individual presynapses were segmented out to create a separate ROI. This ROI was then applied to both the ATG9A and tubulin channels to measure the ATG9A signal intensity per presynapse volume. To obtain the ATG9A intensity specifically in axons (minus presynapse), the ATG9A signal intensity per presynapse volume was subtracted from the ATG9A intensity per axon volume.

#### Quantification of cytochrome c leakage and mitochondrial density

Three separate analysis steps were conducted for the cytochrome *c* (cyt *c*) protruding mitochondria study. First, the percentage of mitochondrial volume lacking cyt *c* in specific neuronal compartments was measured and raw max-projected images were analysed using ImageJ. Somas and presynapses were manually segmented using tubulin staining for morphological distinction, with presynapses identified using the presynaptic marker Synapsin I. The volume occupied by TOM20 and cyt *c* in axons and presynapses were quantified using the same thresholding and “Analyse Particles” methods and compartment segmentation method (i.e. separating somas, axons and presynapses) as described above. The percentage of mitochondrial volume lacking cyt *c* were calculated by the following mathematical formula (*Total TOM*20 *volume* – *Total cyt c volume*) / *Total TOM*20 *volume*) *x* 100%). Second, the percentage of presynapses with mitochondria lacking cyt *c* (i.e. damaged mitochondria) was calculated. The number of total presynapses was manually counted in the segmented images. Each presynapse was examined using a 6.5X zoom image to determine whether any mitochondria were lacking cyt *c*. Third, the average density of total mitochondrial volume per presynapse were measured by dividing the TOM20 volume by the volume occupied by the tubulin in the presynapse segmented images.

### Quantification and Statistical Analysis

For Western blot analysis, band intensities were quantified using ImageLab ver. 6.1 (BioRad). For Image analysis, signal intensities were quantified using ImageJ/Fiji ver.1.54f. Statistical analyses were conducted using GraphPad Prism 10, where significance was determined from three independent experiments using either one-way or two-way ANOVA as specified in the figure legends, with a confidence interval of 95%. Error bars represent either the mean ± standard deviation (SD) or standard error of the mean (SEM), as indicated in the figure legends.

## KEY RESOURCES TABLE

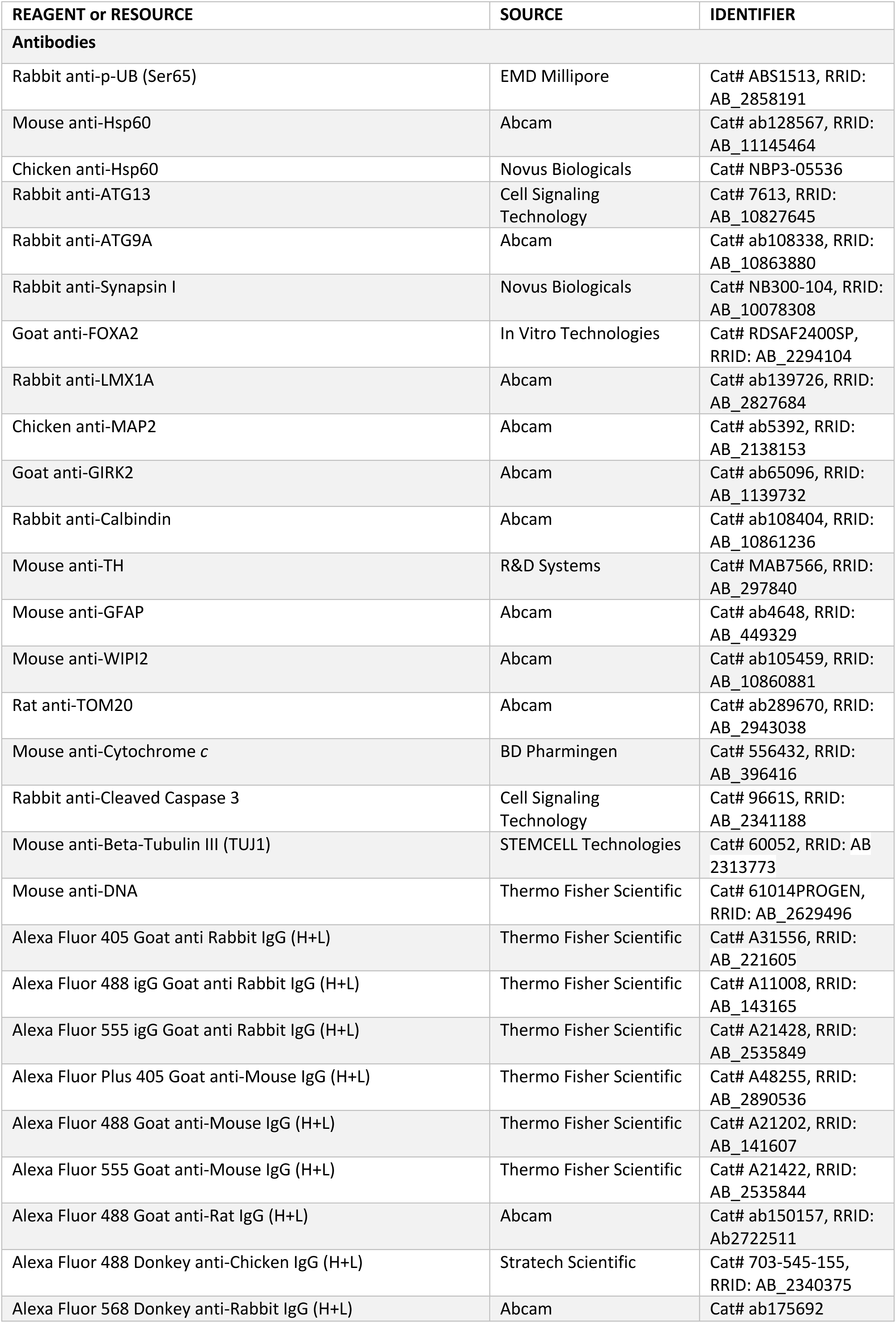
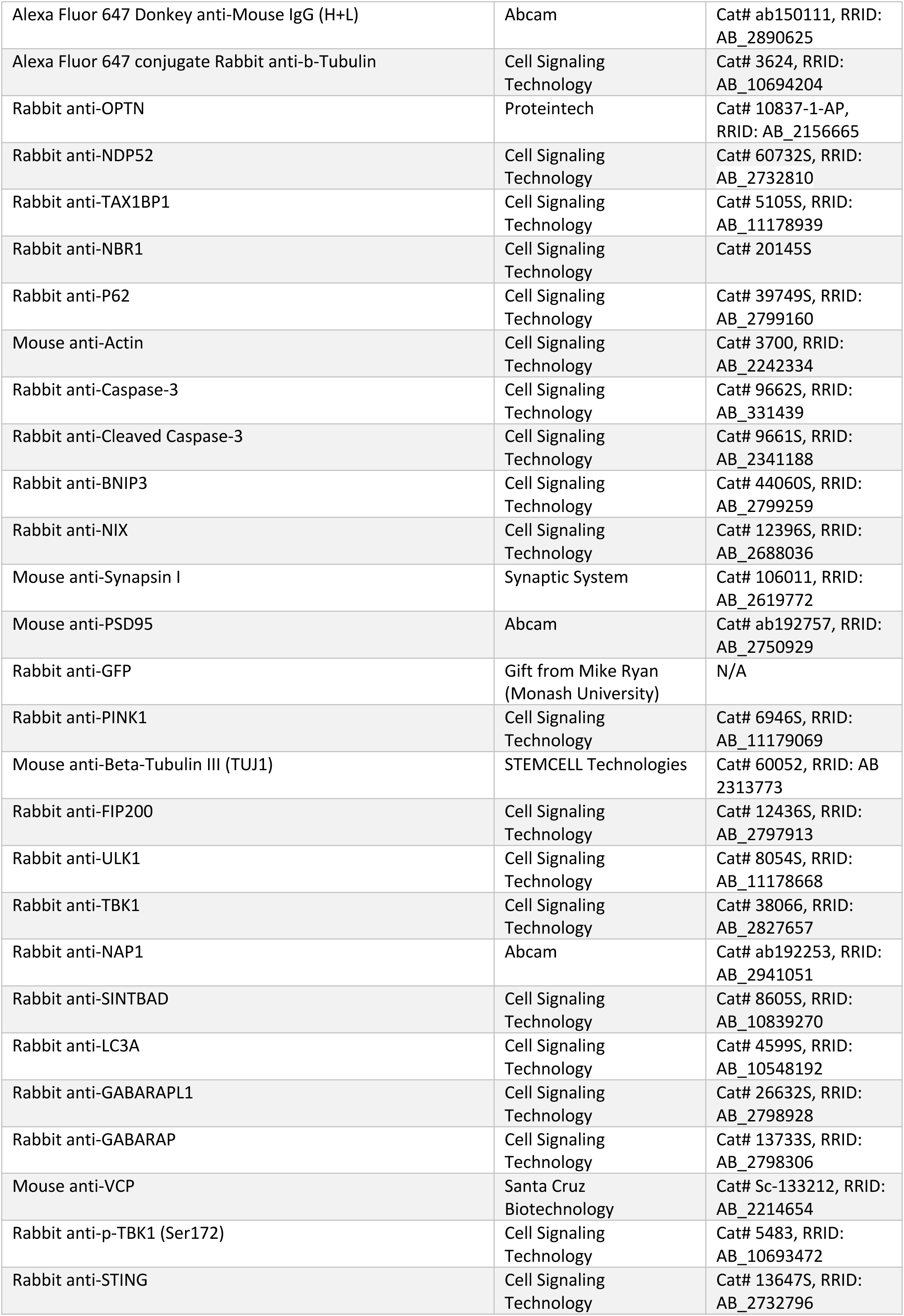
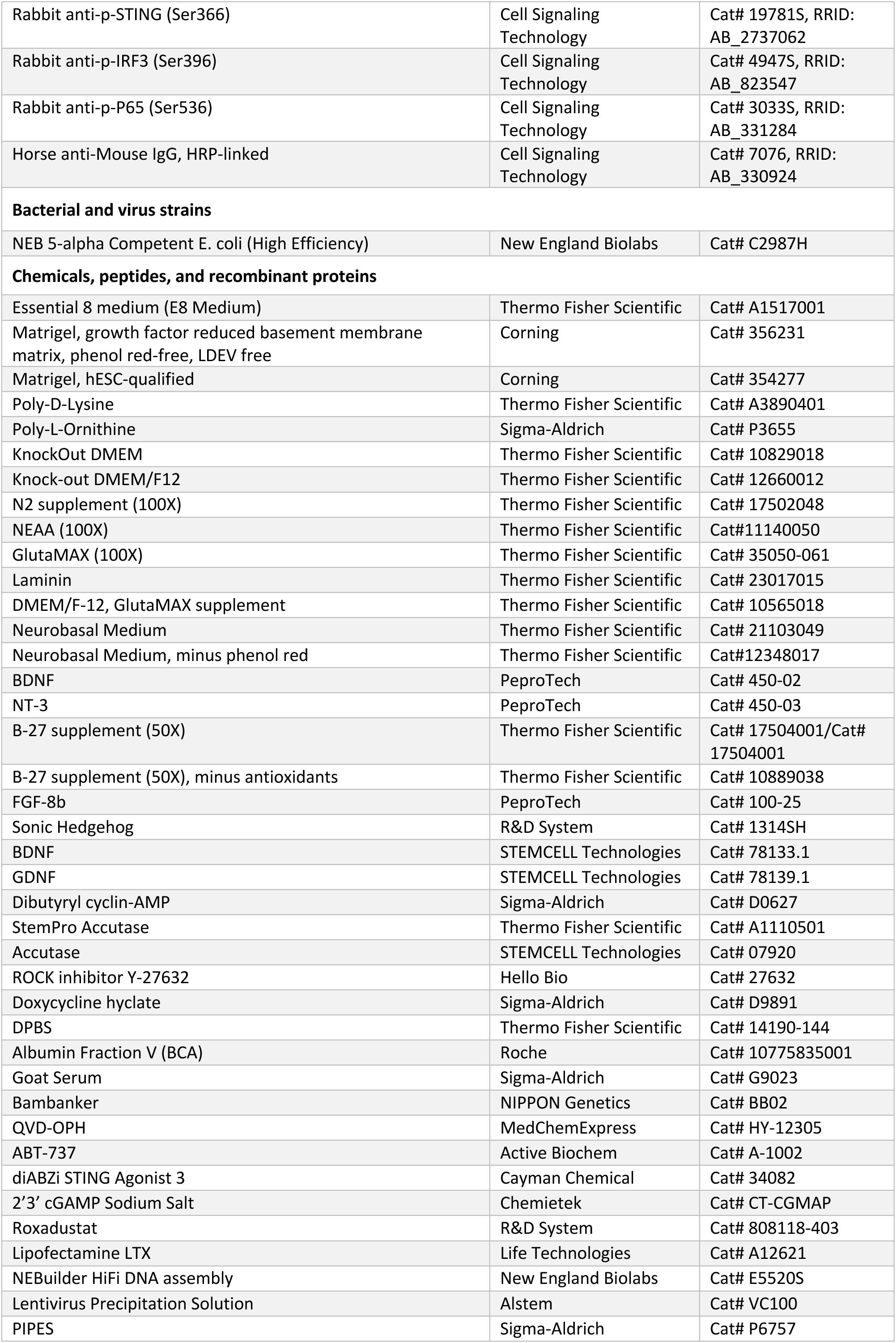
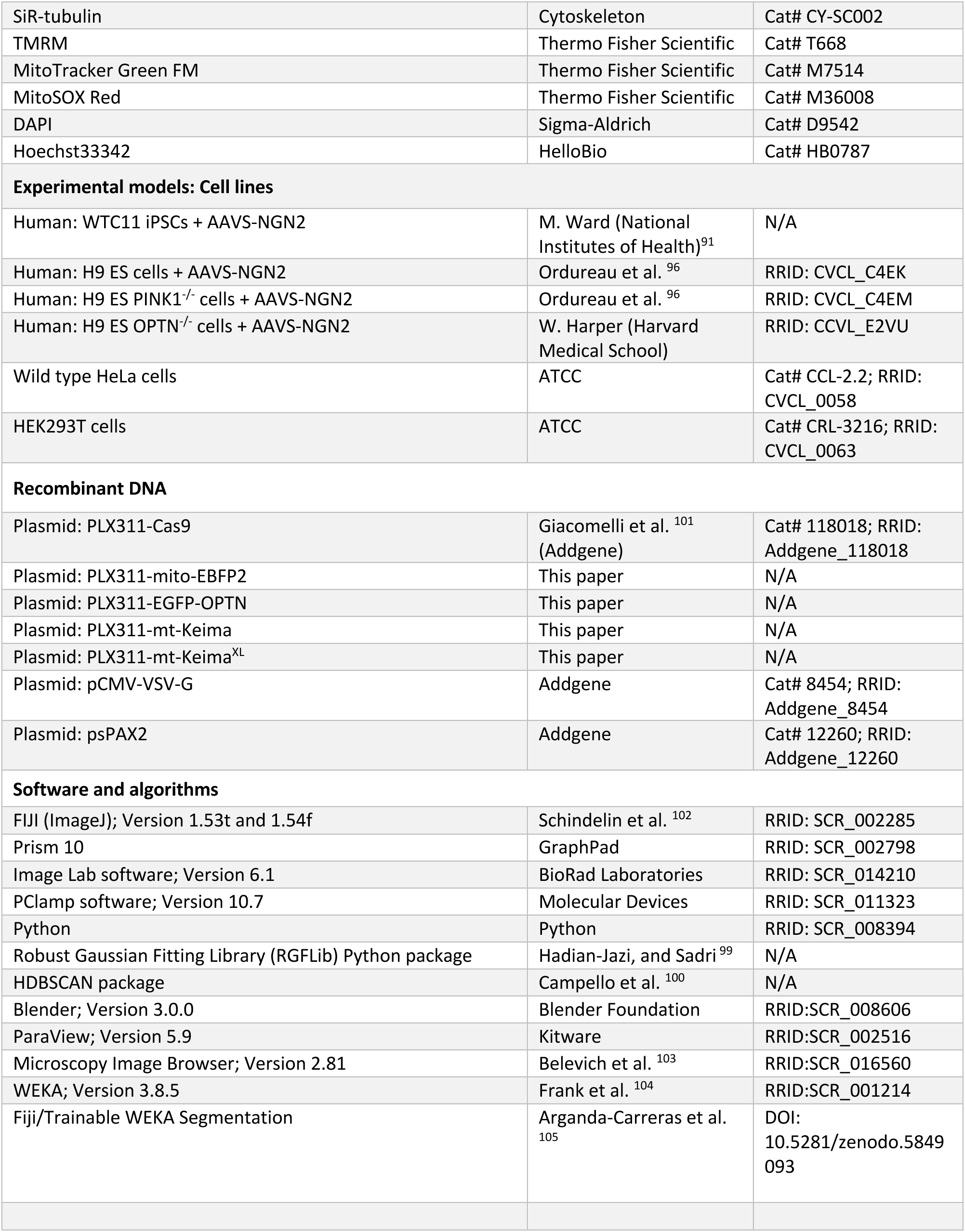

## Acknowledgments

We thank members of the Lazarou Lab and others in Bardy Lab (South Australian Health and Medical Research Institute and Flinders University), Dewson Lab (Walter and Eliza Hall Institute of Medical Research) and Aligning Science Across Parkinson’s (ASAP) Team mito911 for their advice and discussions. We thank Jiuchun Zhang (Harvard Medical School) and Wade Harper (Harvard Medical School) for sharing the WT, PINK1 KO and OPTN KO H9 ESC lines, Michael Ward (National Institutes of Health) for sharing the WTC11 iPSC line and Mike Ryan (Monash Biomedicine Discovery Institute, Monash University) for sharing anti-GFP antibodies. We also thank the WEHI Centre for Dynamic Imaging, the Ramaciotti Centre for Cryo-Electron Microscopy, the WEHI Research Computing Platform, the Monash Micro Imaging Platform, and the WEHI Flow Cytometry Facility for their excellent technical assistance. The summarizing schematic was generated with BioRender. This work was supported by the National Health and Medical Research Council (NHMRC) (GNT1106471 to M.L.), the Australian Research Council Discovery Project (DP200100347 to M.L.) and the Rebecca Cooper Foundation Fellowship (RC20241396 to M.L.), the combined efforts of the Michael J Fox Foundation for Parkinson’s Research (MJFF) and Aligning Science Across Parkinson’s (ASAP) initiative (ASAP-000350 to M.L.); an Australian Research Council Future Fellowship (FT230100138 to C.B.), the Michael J Fox Foundation (to C.B.), Shake It Up Foundation (to C.B.), The Hospital Research Foundation (to C.B.), The Grosset Gaya Fund (to C.B.); NHMRC (GNT2004446 to G.D.), Bodhi Education Fund (to G.D.); The Flinders University Research Scholarship (to P.M.).

## Author contributions

W.K.L. and M.L. conceived the projects. W.K.L., C.B. and M.L. designed the experiments. W.K.L., R.S.J.L., B.M., P.M., C.K., G.K., L.U., T.N.N. and M.S. performed the experiments. W.K.L., R.S.J.L., B.M., P.M., M.H-J. performed experimental data analysis. T.L.S., M.F.S., G.D., A.I.F. contributed to editing the manuscript and conceptual input. W.K.L. and M.L., wrote the manuscript, and all authors contributed to preparing and editing the manuscript.

## References

1. Dikic, I., and Elazar, Z. (2018). Mechanism and medical implications of mammalian autophagy. Nat Rev Mol Cell Biol 19, 349–364. 10.1038/s41580-018-0003-4.

2. Melia, T.J., Lystad, A.H., and Simonsen, A. (2020). Autophagosome biogenesis: From membrane growth to closure. J Cell Biol 219. 10.1083/jcb.202002085.

3. Mercer, T.J., Gubas, A., and Tooze, S.A. (2018). A molecular perspective of mammalian autophagosome biogenesis. J Biol Chem 293, 5386–5395. 10.1074/jbc.R117.810366.

4. Osawa, T., Kotani, T., Kawaoka, T., Hirata, E., Suzuki, K., Nakatogawa, H., Ohsumi, Y., and Noda, N.N. (2019). Atg2 mediates direct lipid transfer between membranes for autophagosome formation. Nat Struct Mol Biol 26, 281–288. 10.1038/s41594-019-0203-4.

5. Sawa-Makarska, J., Baumann, V., Coudevylle, N., von Bulow, S., Nogellova, V., Abert, C., Schuschnig, M., Graef, M., Hummer, G., and Martens, S. (2020). Reconstitution of autophagosome nucleation defines Atg9 vesicles as seeds for membrane formation. Science 369. 10.1126/science.aaz7714.

6. Orsi, A., Razi, M., Dooley, H.C., Robinson, D., Weston, A.E., Collinson, L.M., and Tooze, S.A. (2012). Dynamic and transient interactions of Atg9 with autophagosomes, but not membrane integration, are required for autophagy. Mol Biol Cell 23, 1860–1873. 10.1091/mbc.E11-09-0746.

7. Maeda, S., Yamamoto, H., Kinch, L.N., Garza, C.M., Takahashi, S., Otomo, C., Grishin, N.V., Forli, S., Mizushima, N., and Otomo, T. (2020). Structure, lipid scrambling activity and role in autophagosome formation of ATG9A. Nat Struct Mol Biol 27, 1194–1201. 10.1038/s41594-020-00520-2.

8. van Vliet, A.R., Chiduza, G.N., Maslen, S.L., Pye, V.E., Joshi, D., De Tito, S., Jefferies, H.B.J., Christodoulou, E., Roustan, C., Punch, E., et al. (2022). ATG9A and ATG2A form a heteromeric complex essential for autophagosome formation. Mol Cell 82, 4324–4339 e4328. 10.1016/j.molcel.2022.10.017.

9. Valverde, D.P., Yu, S., Boggavarapu, V., Kumar, N., Lees, J.A., Walz, T., Reinisch, K.M., and Melia, T.J. (2019). ATG2 transports lipids to promote autophagosome biogenesis. J Cell Biol 218, 1787–1798. 10.1083/jcb.201811139.

10. Nguyen, T.N., Sawa-Makarska, J., Khuu, G., Lam, W.K., Adriaenssens, E., Fracchiolla, D., Shoebridge, S., Bernklau, D., Padman, B.S., Skulsuppaisarn, M., et al. (2023). Unconventional initiation of PINK1/Parkin mitophagy by Optineurin. Mol Cell 83, 1693–1709 e1699. 10.1016/j.molcel.2023.04.021.

11. Kotani, T., Kirisako, H., Koizumi, M., Ohsumi, Y., and Nakatogawa, H. (2018). The Atg2-Atg18 complex tethers pre-autophagosomal membranes to the endoplasmic reticulum for autophagosome formation. Proc Natl Acad Sci U S A 115, 10363–10368. 10.1073/pnas.1806727115.

12. Axe, E.L., Walker, S.A., Manifava, M., Chandra, P., Roderick, H.L., Habermann, A., Griffiths, G., and Ktistakis, N.T. (2008). Autophagosome formation from membrane compartments enriched in phosphatidylinositol 3-phosphate and dynamically connected to the endoplasmic reticulum. J Cell Biol 182, 685–701. 10.1083/jcb.200803137.

13. Hayashi-Nishino, M., Fujita, N., Noda, T., Yamaguchi, A., Yoshimori, T., and Yamamoto, A. (2009). A subdomain of the endoplasmic reticulum forms a cradle for autophagosome formation. Nat Cell Biol 11, 1433–1437. 10.1038/ncb1991.

14. Uoselis, L., Nguyen, T.N., and Lazarou, M. (2023). Mitochondrial degradation: Mitophagy and beyond. Mol Cell 83, 3404–3420. 10.1016/j.molcel.2023.08.021.

15. Ordureau, A., Kraus, F., Zhang, J., An, H., Park, S., Ahfeldt, T., Paulo, J.A., and Harper, J.W. (2021). Temporal proteomics during neurogenesis reveals large-scale proteome and organelle remodeling via selective autophagy. Mol Cell 81, 5082–5098 e5011. 10.1016/j.molcel.2021.10.001.

16. Sandoval, H., Thiagarajan, P., Dasgupta, S.K., Schumacher, A., Prchal, J.T., Chen, M., and Wang, J. (2008). Essential role for Nix in autophagic maturation of erythroid cells. Nature 454, 232–235. 10.1038/nature07006.

17. Zhu, Y., Massen, S., Terenzio, M., Lang, V., Chen-Lindner, S., Eils, R., Novak, I., Dikic, I., Hamacher-Brady, A., and Brady, N.R. (2013). Modulation of serines 17 and 24 in the LC3-interacting region of Bnip3 determines pro-survival mitophagy versus apoptosis. J Biol Chem 288, 1099–1113. 10.1074/jbc.M112.399345.

18. Rogov, V.V., Suzuki, H., Marinkovic, M., Lang, V., Kato, R., Kawasaki, M., Buljubasic, M., Sprung, M., Rogova, N., Wakatsuki, S., et al. (2017). Phosphorylation of the mitochondrial autophagy receptor Nix enhances its interaction with LC3 proteins. Sci Rep 7, 1131. 10.1038/s41598-017-01258-6.

19. Bunker, E.N., Le Guerroue, F., Wang, C., Strub, M.P., Werner, A., Tjandra, N., and Youle, R.J. (2023). Nix interacts with WIPI2 to induce mitophagy. EMBO J 42, e113491. 10.15252/embj.2023113491.

20. Slobodkin, M.R., and Elazar, Z. (2013). The Atg8 family: multifunctional ubiquitin-like key regulators of autophagy. Essays Biochem 55, 51–64. 10.1042/bse0550051.

21. Cao, Y., Zheng, J., Wan, H., Sun, Y., Fu, S., Liu, S., He, B., Cai, G., Cao, Y., Huang, H., et al. (2023). A mitochondrial SCF-FBXL4 ubiquitin E3 ligase complex degrades BNIP3 and NIX to restrain mitophagy and prevent mitochondrial disease. EMBO J 42, e113033. 10.15252/embj.2022113033.

22. Elcocks, H., Brazel, A.J., McCarron, K.R., Kaulich, M., Husnjak, K., Mortiboys, H., Clague, M.J., and Urbe, S. (2023). FBXL4 ubiquitin ligase deficiency promotes mitophagy by elevating NIX levels. EMBO J 42, e112799. 10.15252/embj.2022112799.

23. Nguyen-Dien, G.T., Kozul, K.L., Cui, Y., Townsend, B., Kulkarni, P.G., Ooi, S.S., Marzio, A., Carrodus, N., Zuryn, S., Pagano, M., et al. (2023). FBXL4 suppresses mitophagy by restricting the accumulation of NIX and BNIP3 mitophagy receptors. EMBO J 42, e112767. 10.15252/embj.2022112767.

24. Sun, Y., Cao, Y., Wan, H., Memetimin, A., Cao, Y., Li, L., Wu, C., Wang, M., Chen, S., Li, Q., et al. (2024). A mitophagy sensor PPTC7 controls BNIP3 and NIX degradation to regulate mitochondrial mass. Mol Cell 84, 327–344 e329. 10.1016/j.molcel.2023.11.038.

25. Guo, K., Searfoss, G., Krolikowski, D., Pagnoni, M., Franks, C., Clark, K., Yu, K.T., Jaye, M., and Ivashchenko, Y. (2001). Hypoxia induces the expression of the pro-apoptotic gene BNIP3. Cell Death Differ 8, 367–376. 10.1038/sj.cdd.4400810.

26. Bellot, G., Garcia-Medina, R., Gounon, P., Chiche, J., Roux, D., Pouyssegur, J., and Mazure, N.M. (2009). Hypoxia-induced autophagy is mediated through hypoxia-inducible factor induction of BNIP3 and BNIP3L via their BH3 domains. Mol Cell Biol 29, 2570–2581. 10.1128/MCB.00166-09.

27. Allen, G.F., Toth, R., James, J., and Ganley, I.G. (2013). Loss of iron triggers PINK1/Parkin-independent mitophagy. EMBO Rep 14, 1127–1135. 10.1038/embor.2013.168.

28. Kitada, T., Asakawa, S., Hattori, N., Matsumine, H., Yamamura, Y., Minoshima, S., Yokochi, M., Mizuno, Y., and Shimizu, N. (1998). Mutations in the parkin gene cause autosomal recessive juvenile parkinsonism. Nature 392, 605–608. 10.1038/33416.

29. Valente, E.M., Bentivoglio, A.R., Dixon, P.H., Ferraris, A., Ialongo, T., Frontali, M., Albanese, A., and Wood, N.W. (2001). Localization of a novel locus for autosomal recessive early-onset parkinsonism, PARK6, on human chromosome 1p35-p36. Am J Hum Genet 68, 895–900. 10.1086/319522.

30. Fang, E.F., Hou, Y., Palikaras, K., Adriaanse, B.A., Kerr, J.S., Yang, B., Lautrup, S., Hasan-Olive, M.M., Caponio, D., Dan, X., et al. (2019). Mitophagy inhibits amyloid-beta and tau pathology and reverses cognitive deficits in models of Alzheimer’s disease. Nat Neurosci 22, 401–412. 10.1038/s41593-018-0332-9.

31. Yi, J., Wang, H.L., Lu, G., Zhang, H., Wang, L., Li, Z.Y., Wang, L., Wu, Y., Xia, D., Fang, E.F., and Shen, H.M. (2024). Spautin-1 promotes PINK1-PRKN-dependent mitophagy and improves associative learning capability in an alzheimer disease animal model. Autophagy, 1-22. 10.1080/15548627.2024.2383145.

32. Evans, C.S., and Holzbaur, E.L.F. (2019). Autophagy and mitophagy in ALS. Neurobiol Dis 122, 35–40. 10.1016/j.nbd.2018.07.005.

33. Burke, R.E., and O’Malley, K. (2013). Axon degeneration in Parkinson’s disease. Exp Neurol 246, 72–83. 10.1016/j.expneurol.2012.01.011.

34. Padmanabhan, P., Kneynsberg, A., and Gotz, J. (2021). Super-resolution microscopy: a closer look at synaptic dysfunction in Alzheimer disease. Nat Rev Neurosci 22, 723–740. 10.1038/s41583-021-00531-y.

35. Kordower, J.H., Olanow, C.W., Dodiya, H.B., Chu, Y., Beach, T.G., Adler, C.H., Halliday, G.M., and Bartus, R.T. (2013). Disease duration and the integrity of the nigrostriatal system in Parkinson’s disease. Brain 136, 2419–2431. 10.1093/brain/awt192.

36. Raff, M.C., Whitmore, A.V., and Finn, J.T. (2002). Axonal self-destruction and neurodegeneration. Science 296, 868–871. 10.1126/science.1068613.

37. Nikoletopoulou, V., Papandreou, M.E., and Tavernarakis, N. (2015). Autophagy in the physiology and pathology of the central nervous system. Cell Death Differ 22, 398–407. 10.1038/cdd.2014.204.

38. Perez, F.A., and Palmiter, R.D. (2005). Parkin-deficient mice are not a robust model of parkinsonism. Proc Natl Acad Sci U S A 102, 2174–2179. 10.1073/pnas.0409598102.

39. Kitada, T., Pisani, A., Porter, D.R., Yamaguchi, H., Tscherter, A., Martella, G., Bonsi, P., Zhang, C., Pothos, E.N., and Shen, J. (2007). Impaired dopamine release and synaptic plasticity in the striatum of PINK1-deficient mice. Proc Natl Acad Sci U S A 104, 11441–11446. 10.1073/pnas.0702717104.

40. de Haas, R., Heltzel, L., Tax, D., van den Broek, P., Steenbreker, H., Verheij, M.M.M., Russel, F.G.M., Orr, A.L., Nakamura, K., and Smeitink, J.A.M. (2019). To be or not to be pink(1): contradictory findings in an animal model for Parkinson’s disease. Brain Commun 1, fcz016. 10.1093/braincomms/fcz016.

41. Goldsmith, J., Ordureau, A., Boecker CA, C.A., Arany, M., Harper, J.W., and Holzbaur, E.L. (2023). Proteomics analysis of autophagy cargos reveals distinct adaptations in PINK1 and LRRK2 models of Parkinson disease. bioRxiv, 2022.2010.2003.510717. 10.1101/2022.10.03.510717.

42. Evans, C.S., and Holzbaur, E.L. (2020). Degradation of engulfed mitochondria is rate-limiting in Optineurin-mediated mitophagy in neurons. Elife 9. 10.7554/eLife.50260.

43. Fernandopulle, M.S., Prestil, R., Grunseich, C., Wang, C., Gan, L., and Ward, M.E. (2018). Transcription Factor-Mediated Differentiation of Human iPSCs into Neurons. Curr Protoc Cell Biol 79, e51. 10.1002/cpcb.51.

44. Cheng, X.T., Zhou, B., Lin, M.Y., Cai, Q., and Sheng, Z.H. (2015). Axonal autophagosomes recruit dynein for retrograde transport through fusion with late endosomes. J Cell Biol 209, 377–386. 10.1083/jcb.201412046.

45. Hollenbeck, P.J. (1993). Products of endocytosis and autophagy are retrieved from axons by regulated retrograde organelle transport. J Cell Biol 121, 305–315. 10.1083/jcb.121.2.305.

46. Maday, S., Wallace, K.E., and Holzbaur, E.L. (2012). Autophagosomes initiate distally and mature during transport toward the cell soma in primary neurons. J Cell Biol 196, 407–417. 10.1083/jcb.201106120.

47. Lie, P.P.Y., Yang, D.S., Stavrides, P., Goulbourne, C.N., Zheng, P., Mohan, P.S., Cataldo, A.M., and Nixon, R.A. (2021). Post-Golgi carriers, not lysosomes, confer lysosomal properties to pre-degradative organelles in normal and dystrophic axons. Cell Rep 35, 109034. 10.1016/j.celrep.2021.109034.

48. Kane, L.A., Lazarou, M., Fogel, A.I., Li, Y., Yamano, K., Sarraf, S.A., Banerjee, S., and Youle, R.J. (2014). PINK1 phosphorylates ubiquitin to activate Parkin E3 ubiquitin ligase activity. J Cell Biol 205, 143–153. 10.1083/jcb.201402104.

49. Kazlauskaite, A., Kondapalli, C., Gourlay, R., Campbell, D.G., Ritorto, M.S., Hofmann, K., Alessi, D.R., Knebel, A., Trost, M., and Muqit, M.M. (2014). Parkin is activated by PINK1-dependent phosphorylation of ubiquitin at Ser65. Biochem J 460, 127–139. 10.1042/BJ20140334.

50. Koyano, F., Okatsu, K., Kosako, H., Tamura, Y., Go, E., Kimura, M., Kimura, Y., Tsuchiya, H., Yoshihara, H., Hirokawa, T., et al. (2014). Ubiquitin is phosphorylated by PINK1 to activate parkin. Nature 510, 162–166. 10.1038/nature13392.

51. Ordureau, A., Heo, J.M., Duda, D.M., Paulo, J.A., Olszewski, J.L., Yanishevski, D., Rinehart, J., Schulman, B.A., and Harper, J.W. (2015). Defining roles of PARKIN and ubiquitin phosphorylation by PINK1 in mitochondrial quality control using a ubiquitin replacement strategy. Proc Natl Acad Sci U S A 112, 6637–6642. 10.1073/pnas.1506593112.

52. Okatsu, K., Koyano, F., Kimura, M., Kosako, H., Saeki, Y., Tanaka, K., and Matsuda, N. (2015). Phosphorylated ubiquitin chain is the genuine Parkin receptor. J Cell Biol 209, 111–128. 10.1083/jcb.201410050.

53. Wauer, T., Simicek, M., Schubert, A., and Komander, D. (2015). Mechanism of phospho-ubiquitin-induced PARKIN activation. Nature 524, 370–374. 10.1038/nature14879.

54. Harbauer, A.B., Hees, J.T., Wanderoy, S., Segura, I., Gibbs, W., Cheng, Y., Ordonez, M., Cai, Z., Cartoni, R., Ashrafi, G., et al. (2022). Neuronal mitochondria transport Pink1 mRNA via synaptojanin 2 to support local mitophagy. Neuron 110, 1516–1531 e1519. 10.1016/j.neuron.2022.01.035.

55. Yang, S., Park, D., Manning, L., Hill, S.E., Cao, M., Xuan, Z., Gonzalez, I., Dong, Y., Clark, B., Shao, L., et al. (2022). Presynaptic autophagy is coupled to the synaptic vesicle cycle via ATG-9. Neuron 110, 824–840 e810. 10.1016/j.neuron.2021.12.031.

56. Lazarou, M., Sliter, D.A., Kane, L.A., Sarraf, S.A., Wang, C., Burman, J.L., Sideris, D.P., Fogel, A.I., and Youle, R.J. (2015). The ubiquitin kinase PINK1 recruits autophagy receptors to induce mitophagy. Nature 524, 309–314. 10.1038/nature14893.

57. Bardy, C., van den Hurk, M., Eames, T., Marchand, C., Hernandez, R.V., Kellogg, M., Gorris, M., Galet, B., Palomares, V., Brown, J., et al. (2015). Neuronal medium that supports basic synaptic functions and activity of human neurons in vitro. Proc Natl Acad Sci U S A 112, E2725–2734. 10.1073/pnas.1504393112.

58. Zabolocki, M., McCormack, K., van den Hurk, M., Milky, B., Shoubridge, A.P., Adams, R., Tran, J., Mahadevan-Jansen, A., Reineck, P., Thomas, J., et al. (2020). BrainPhys neuronal medium optimized for imaging and optogenetics in vitro. Nat Commun 11, 5550. 10.1038/s41467-020-19275-x.

59. Milky, B., Zabolocki, M., Al-Bataineh, S.A., van den Hurk, M., Greenberg, Z., Turner, L., Mazzachi, P., Williams, A., Illeperuma, I., Adams, R., et al. (2022). Long-term adherence of human brain cells in vitro is enhanced by charged amine-based plasma polymer coatings. Stem Cell Reports 17, 489–506. 10.1016/j.stemcr.2022.01.013.

60. Tran, J., Anastacio, H., and Bardy, C. (2020). Genetic predispositions of Parkinson’s disease revealed in patient-derived brain cells. NPJ Parkinsons Dis 6, 8. 10.1038/s41531-020-0110-8.

61. Shapson-Coe, A., Januszewski, M., Berger, D.R., Pope, A., Wu, Y., Blakely, T., Schalek, R.L., Li, P.H., Wang, S., Maitin-Shepard, J., et al. (2024). A petavoxel fragment of human cerebral cortex reconstructed at nanoscale resolution. Science 384, eadk4858. 10.1126/science.adk4858.

62. Nguyen, T.N., Padman, B.S., Zellner, S., Khuu, G., Uoselis, L., Lam, W.K., Skulsuppaisarn, M., Lindblom, R.S.J., Watts, E.M., Behrends, C., and Lazarou, M. (2021). ATG4 family proteins drive phagophore growth independently of the LC3/GABARAP lipidation system. Mol Cell 81, 2013–2030 e2019. 10.1016/j.molcel.2021.03.001.

63. Slee, E.A., Harte, M.T., Kluck, R.M., Wolf, B.B., Casiano, C.A., Newmeyer, D.D., Wang, H.G., Reed, J.C., Nicholson, D.W., Alnemri, E.S., et al. (1999). Ordering the cytochrome c-initiated caspase cascade: hierarchical activation of caspases-2, -3, -6, -7, -8, and -10 in a caspase-9-dependent manner. J Cell Biol 144, 281–292. 10.1083/jcb.144.2.281.

64. Janicke, R.U., Sprengart, M.L., Wati, M.R., and Porter, A.G. (1998). Caspase-3 is required for DNA fragmentation and morphological changes associated with apoptosis. J Biol Chem 273, 9357–9360. 10.1074/jbc.273.16.9357.

65. Kluck, R.M., Bossy-Wetzel, E., Green, D.R., and Newmeyer, D.D. (1997). The release of cytochrome c from mitochondria: a primary site for Bcl-2 regulation of apoptosis. Science 275, 1132–1136. 10.1126/science.275.5303.1132.

66. Jiang, X., and Wang, X. (2004). Cytochrome C-mediated apoptosis. Annu Rev Biochem 73, 87–106. 10.1146/annurev.biochem.73.011303.073706.

67. White, M.J., McArthur, K., Metcalf, D., Lane, R.M., Cambier, J.C., Herold, M.J., van Delft, M.F., Bedoui, S., Lessene, G., Ritchie, M.E., et al. (2014). Apoptotic caspases suppress mtDNA-induced STING-mediated type I IFN production. Cell 159, 1549–1562. 10.1016/j.cell.2014.11.036.

68. McArthur, K., Whitehead, L.W., Heddleston, J.M., Li, L., Padman, B.S., Oorschot, V., Geoghegan, N.D., Chappaz, S., Davidson, S., San Chin, H., et al. (2018). BAK/BAX macropores facilitate mitochondrial herniation and mtDNA efflux during apoptosis. Science 359. 10.1126/science.aao6047.

69. Riley, J.S., Quarato, G., Cloix, C., Lopez, J., O’Prey, J., Pearson, M., Chapman, J., Sesaki, H., Carlin, L.M., Passos, J.F., et al. (2018). Mitochondrial inner membrane permeabilisation enables mtDNA release during apoptosis. EMBO J 37. 10.15252/embj.201899238.

70. Rongvaux, A., Jackson, R., Harman, C.C., Li, T., West, A.P., de Zoete, M.R., Wu, Y., Yordy, B., Lakhani, S.A., Kuan, C.Y., et al. (2014). Apoptotic caspases prevent the induction of type I interferons by mitochondrial DNA. Cell 159, 1563–1577. 10.1016/j.cell.2014.11.037.

71. Zhao, J.F., Rodger, C.E., Allen, G.F.G., Weidlich, S., and Ganley, I.G. (2020). HIF1alpha-dependent mitophagy facilitates cardiomyoblast differentiation. Cell Stress 4, 99–113. 10.15698/cst2020.05.220.

72. Ivan, M., Kondo, K., Yang, H., Kim, W., Valiando, J., Ohh, M., Salic, A., Asara, J.M., Lane, W.S., and Kaelin, W.G., Jr. (2001). HIFalpha targeted for VHL-mediated destruction by proline hydroxylation: implications for O2 sensing. Science 292, 464–468. 10.1126/science.1059817.

73. Jaakkola, P., Mole, D.R., Tian, Y.M., Wilson, M.I., Gielbert, J., Gaskell, S.J., von Kriegsheim, A., Hebestreit, H.F., Mukherji, M., Schofield, C.J., et al. (2001). Targeting of HIF-alpha to the von Hippel-Lindau ubiquitylation complex by O2-regulated prolyl hydroxylation. Science 292, 468–472. 10.1126/science.1059796.

74. Haase, V.H. (2021). Hypoxia-inducible factor-prolyl hydroxylase inhibitors in the treatment of anemia of chronic kidney disease. Kidney Int Suppl (2011) 11, 8–25. 10.1016/j.kisu.2020.12.002.

75. Singh, A.K., Carroll, K., McMurray, J.J.V., Solomon, S., Jha, V., Johansen, K.L., Lopes, R.D., Macdougall, I.C., Obrador, G.T., Waikar, S.S., et al. (2021). Daprodustat for the Treatment of Anemia in Patients Not Undergoing Dialysis. N Engl J Med 385, 2313–2324. 10.1056/NEJMoa2113380.

76. Hamanaka, R.B., and Chandel, N.S. (2009). Mitochondrial reactive oxygen species regulate hypoxic signaling. Curr Opin Cell Biol 21, 894–899. 10.1016/j.ceb.2009.08.005.

77. Ashrafi, G., Schlehe, J.S., LaVoie, M.J., and Schwarz, T.L. (2014). Mitophagy of damaged mitochondria occurs locally in distal neuronal axons and requires PINK1 and Parkin. J Cell Biol 206, 655–670. 10.1083/jcb.201401070.

78. Kang, J.S., Tian, J.H., Pan, P.Y., Zald, P., Li, C., Deng, C., and Sheng, Z.H. (2008). Docking of axonal mitochondria by syntaphilin controls their mobility and affects short-term facilitation. Cell 132, 137–148. 10.1016/j.cell.2007.11.024.

79. Courchet, J., Lewis, T.L., Jr., Lee, S., Courchet, V., Liou, D.Y., Aizawa, S., and Polleux, F. (2013). Terminal axon branching is regulated by the LKB1-NUAK1 kinase pathway via presynaptic mitochondrial capture. Cell 153, 1510–1525. 10.1016/j.cell.2013.05.021.

80. Wang, X., Winter, D., Ashrafi, G., Schlehe, J., Wong, Y.L., Selkoe, D., Rice, S., Steen, J., LaVoie, M.J., and Schwarz, T.L. (2011). PINK1 and Parkin target Miro for phosphorylation and degradation to arrest mitochondrial motility. Cell 147, 893–906. 10.1016/j.cell.2011.10.018.

81. Wang, X., and Schwarz, T.L. (2009). The mechanism of Ca2+ -dependent regulation of kinesin-mediated mitochondrial motility. Cell 136, 163–174. 10.1016/j.cell.2008.11.046.

82. Park, D., Wu, Y., Wang, X., Gowrishankar, S., Baublis, A., and De Camilli, P. (2023). Synaptic vesicle proteins and ATG9A self-organize in distinct vesicle phases within synapsin condensates. Nat Commun 14, 455. 10.1038/s41467-023-36081-3.

83. Schoenmann, Z., Assa-Kunik, E., Tiomny, S., Minis, A., Haklai-Topper, L., Arama, E., and Yaron, A. (2010). Axonal degeneration is regulated by the apoptotic machinery or a NAD+-sensitive pathway in insects and mammals. J Neurosci 30, 6375–6386. 10.1523/JNEUROSCI.0922-10.2010.

84. Cowan, C.M., Thai, J., Krajewski, S., Reed, J.C., Nicholson, D.W., Kaufmann, S.H., and Roskams, A.J. (2001). Caspases 3 and 9 send a pro-apoptotic signal from synapse to cell body in olfactory receptor neurons. J Neurosci 21, 7099–7109. 10.1523/JNEUROSCI.21-18-07099.2001.

85. Simon, D.J., Weimer, R.M., McLaughlin, T., Kallop, D., Stanger, K., Yang, J., O’Leary, D.D., Hannoush, R.N., and Tessier-Lavigne, M. (2012). A caspase cascade regulating developmental axon degeneration. J Neurosci 32, 17540–17553. 10.1523/JNEUROSCI.3012-12.2012.

86. Mukherjee, A., and Williams, D.W. (2017). More alive than dead: non-apoptotic roles for caspases in neuronal development, plasticity and disease. Cell Death Differ 24, 1411–1421. 10.1038/cdd.2017.64.

87. Cusack, C.L., Swahari, V., Hampton Henley, W., Michael Ramsey, J., and Deshmukh, M. (2013). Distinct pathways mediate axon degeneration during apoptosis and axon-specific pruning. Nat Commun 4, 1876. 10.1038/ncomms2910.

88. Haider, M.U., Furqan, M., and Mehmood, Q. (2023). Daprodustat: A potential game-changer in renal anemia therapy-A perspective. Front Pharmacol 14, 1249492. 10.3389/fphar.2023.1249492.

89. Fang, T.Z., Sun, Y., Pearce, A.C., Eleuteri, S., Kemp, M., Luckhurst, C.A., Williams, R., Mills, R., Almond, S., Burzynski, L., et al. (2023). Knockout or inhibition of USP30 protects dopaminergic neurons in a Parkinson’s disease mouse model. Nat Commun 14, 7295. 10.1038/s41467-023-42876-1.

90. Miyaoka, Y., Chan, A.H., Judge, L.M., Yoo, J., Huang, M., Nguyen, T.D., Lizarraga, P.P., So, P.L., and Conklin, B.R. (2014). Isolation of single-base genome-edited human iPS cells without antibiotic selection. Nat Methods 11, 291–293. 10.1038/nmeth.2840.

91. Wang, C., Ward, M.E., Chen, R., Liu, K., Tracy, T.E., Chen, X., Xie, M., Sohn, P.D., Ludwig, C., Meyer-Franke, A., et al. (2017). Scalable Production of iPSC-Derived Human Neurons to Identify Tau-Lowering Compounds by High-Content Screening. Stem Cell Reports 9, 1221–1233. 10.1016/j.stemcr.2017.08.019.

92. Tian, R., Gachechiladze, M.A., Ludwig, C.H., Laurie, M.T., Hong, J.Y., Nathaniel, D., Prabhu, A.V., Fernandopulle, M.S., Patel, R., Abshari, M., et al. (2019). CRISPR Interference-Based Platform for Multimodal Genetic Screens in Human iPSC-Derived Neurons. Neuron 104, 239–255 e212. 10.1016/j.neuron.2019.07.014.

93. Ordureau, A., Paulo, J.A., Zhang, W., Ahfeldt, T., Zhang, J., Cohn, E.F., Hou, Z., Heo, J.M., Rubin, L.L., Sidhu, S.S., et al. (2018). Dynamics of PARKIN-Dependent Mitochondrial Ubiquitylation in Induced Neurons and Model Systems Revealed by Digital Snapshot Proteomics. Mol Cell 70, 211–227 e218. 10.1016/j.molcel.2018.03.012.

94. Eapen, V.V., Swarup, S., Hoyer, M.J., Paulo, J.A., and Harper, J.W. (2021). Quantitative proteomics reveals the selectivity of ubiquitin-binding autophagy receptors in the turnover of damaged lysosomes by lysophagy. Elife 10. 10.7554/eLife.72328.

95. Boyer, L.F., Campbell, B., Larkin, S., Mu, Y., and Gage, F.H. (2012). Dopaminergic differentiation of human pluripotent cells. Curr Protoc Stem Cell Biol Chapter 1, Unit1H 6. 10.1002/9780470151808.sc01h06s22.

96. Ordureau, A., Paulo, J.A., Zhang, J., An, H., Swatek, K.N., Cannon, J.R., Wan, Q., Komander, D., and Harper, J.W. (2020). Global Landscape and Dynamics of Parkin and USP30-Dependent Ubiquitylomes in iNeurons during Mitophagic Signaling. Mol Cell 77, 1124–1142 e1110. 10.1016/j.molcel.2019.11.013.

97. Silva, M.T., Guerra, F.C., and Magalhaes, M.M. (1968). The fixative action of uranyl acetate in electron microscopy. Experientia 24, 1074. 10.1007/BF02138757.

98. Walton, J. (1979). Lead aspartate, an en bloc contrast stain particularly useful for ultrastructural enzymology. J Histochem Cytochem 27, 1337–1342. 10.1177/27.10.512319.

99. Hadian-Jazi, M., and Sadri, A. (2023). A Python package based on robust statistical analysis for serial crystallography data processing. Acta Crystallogr D Struct Biol 79, 820–829. 10.1107/S2059798323005855.

100. Campello, R.J.G.B., Moulavi, D., and Sander, J. (2013). Density-Based Clustering Based on Hierarchical Density Estimates. Advances in Knowledge Discovery and Data Mining. 10.1007/978-3-642-37456-2_14.

101. Giacomelli, A.O., Yang, X., Lintner, R.E., McFarland, J.M., Duby, M., Kim, J., Howard, T.P., Takeda, D.Y., Ly, S.H., Kim, E., et al. (2018). Mutational processes shape the landscape of TP53 mutations in human cancer. Nat Genet 50, 1381–1387. 10.1038/s41588-018-0204-y.

102. Schindelin, J., Arganda-Carreras, I., Frise, E., Kaynig, V., Longair, M., Pietzsch, T., Preibisch, S., Rueden, C., Saalfeld, S., Schmid, B., et al. (2012). Fiji: an open-source platform for biological-image analysis. Nat Methods 9, 676–682. 10.1038/nmeth.2019.

103. Belevich, I., Joensuu, M., Kumar, D., Vihinen, H., and Jokitalo, E. (2016). Microscopy Image Browser: A Platform for Segmentation and Analysis of Multidimensional Datasets. PLoS Biol 14, e1002340. 10.1371/journal.pbio.1002340.

104. Frank, E., Hall, M.A., and Witten, I.H. (2016). The WEKA Workbench. Online Appendix for Data Mining: Practical Machine Learning Tools and Techniques, Fourth Edition (Morgan Kaufmann).

105. Arganda-Carreras, I., Kaynig, V., Rueden, C., Eliceiri, K.W., Schindelin, J., Cardona, A., and Sebastian Seung, H. (2017). Trainable Weka Segmentation: a machine learning tool for microscopy pixel classification. Bioinformatics 33, 2424–2426. 10.1093/bioinformatics/btx180.

